# Birds invest wingbeats to keep a steady head and reap the ultimate benefits of flocking

**DOI:** 10.1101/492090

**Authors:** Lucy A. Taylor, Dora Biro, Ben Lambert, James A. Walker, Graham K. Taylor, Steven J. Portugal

## Abstract

Flapping flight is the most energetically demanding form of sustained forwards locomotion that vertebrates perform. Flock dynamics therefore have significant implications for energy expenditure. Despite this, no studies have quantified the biomechanical consequences of flying in a cluster flock relative to flying solo. Here, we compared the flight characteristics of homing pigeons (*Columba livia*) flying solo and in pairs, using high-precision 5 Hz GPS and 200 Hz tri-axial accelerometer biologgers. Paired flight increased route accuracy by ~7%, but, was accompanied by an increase in wingbeat frequency of ~18%. As expected, paired individuals benefitted from improved homing route accuracy, which reduced flight distance by ~7% and time by ~9%. However, realising these navigational gains involved substantial changes in flight kinematics and energetics. Both individuals in a pair increased their wingbeat frequency by *c.*18%, by decreasing the duration of their upstroke. This sharp increase in wingbeat frequency caused just a 3% increase in airspeed, but reduced the oscillatory displacement of the body by ~22%, which we hypothesise relates to an increased requirement for visual stability and manoeuvrability when flocking. Overall, the shorter flight distances and increased wingbeat frequency in a pair resulted in a net increase in the aerodynamic cost of returning home, which we estimate was ~14%. Our results demonstrate that flocking costs have been underestimated by an order of magnitude and force reinterpretation of their mechanistic origin. We show that, for pigeons, two heads are better than one, but keeping a steady head necessitates energetically costly kinematics.

## Introduction

Across the animal kingdom many species travel in groups, from pairs to flocks, shoals, herds and swarms, some containing millions of individuals [1,2]. Indeed, the collective motion of animals produces some of the most spectacular displays of synchronisation and coordination in the world [3]. Commonly cited benefits of collective travel include an improved ability to detect and avoid predators [1,4], enhanced orientational efficiency through the pooling of navigational knowledge [5–8], and energetic efficiencies derived from fluid dynamic interactions [9–13]. Flocking in birds, in particular, has received considerable attention due to the complex aerodynamic interactions that take place between group members [11–14].

Avian flock formations can be categorised as either line formations or cluster formations [15,16]. Line formations, which include the distinctive ‘V’ of many long-distance migrants, are utilised by medium to large-sized birds, such as northern bald ibis (*Geronticus eremita*) and Canada geese (*Branta canadensis*), whereas cluster formations are typically observed in smaller birds, such as homing pigeons (*Columba livia*) and common starlings (*Sturnus vulgaris*), which fly in irregular three-dimensional flocks [11–16]. Birds flying in close cluster flocks in particular are able to move with near perfect synchrony, whilst making rapid directional changes in three dimensions. While birds travelling in V-formation can save energy by flying in aerodynamically optimal positioning within the V [11–13], those species flying in cluster flocks have been shown to incur an additional energetic cost in denser formations [14]. In homing pigeons, for example, a tenfold increase in the spatial density of a flock has been observed to be associated with a modest 0.1 Hz increase in wingbeat frequency, and was presumed to be accompanied by an energetic cost over the seven flights that were observed [14]. Flapping flight is the most energetically demanding form of sustained forwards locomotion that vertebrates perform [17,18], and flock dynamics may therefore have significant implications for individual energy expenditure and lifetime fitness. However, no studies have yet compared the biomechanical consequences of flying in a pair to flying solo, so the energetic impact of this form of flocking is unknown.

To fill this fundamental gap, we recorded the body accelerations associated with every wingbeat of 20 free-flying homing pigeons flying solo and in pairs as they homed from a site 7 km east of their loft (Fig. 1A). The birds were equipped with 5 Hz GPS trackers and 200 Hz tri-axial accelerometer biologgers which allowed us to reconstruct their trajectories and wingbeat patterns during each homeward flight (see Methods; Fig. 1B-F) [19,20]. The experiment consisted of four phases. In Phase 1, each subject first completed 21 successive solo flights (Fig. 1F), the last six of which provided the solo baseline (solo 1). During these six solo flights, the median wingbeat frequency and amplitude (dorsal body displacement) for all birds were 5.48±0.19 Hz (grand mean of the median value of each flight±s.d. of the individual means) and 20.68±1.17 mm, respectively (after accounting for the effects of airspeed, date of release and weather variables including wind support, crosswind, temperature, humidity and air density; Fig. 2A). These wingbeat frequencies are consistent with those measured previously for solo pigeon flights [20,21]. In Phase 2, following the solo releases, birds were released six times from the same - now familiar - site, but in similar-sized pairs. Pairs were assigned based on similarity in body mass and structural size as measured by tarsus length [22]. Body size and mass are strong predictors of preferred flight speeds in birds, both at the intra- and inter-specific levels, with optimal flight speeds usually assumed to be those for which the cost of transport (i.e. energy expenditure per unit distance) is predicted to be at its minimum [18]. Therefore, we hypothesised that for birds of different sizes either one or both birds may have to adjust their wingbeat frequency and/or amplitude to stay together as a pair, which would represent an additional ‘hidden’ compromise cost of flying with another bird. In Phase 3, immediately following completion of the six similar-sized pair releases, each bird was then flown in size-mismatched pairs for a further six flights (different-sized pairs), again from the same site. Finally, in Phase 4, upon completion of the six size-mismatched flights, each bird flew six times solo again (Fig. 2D). We compared the wingbeat characteristics of birds flying in pairs relative to flying solo to determine if pigeons alter their wingbeat characteristics when flying in a pair.

**Figure 1.**
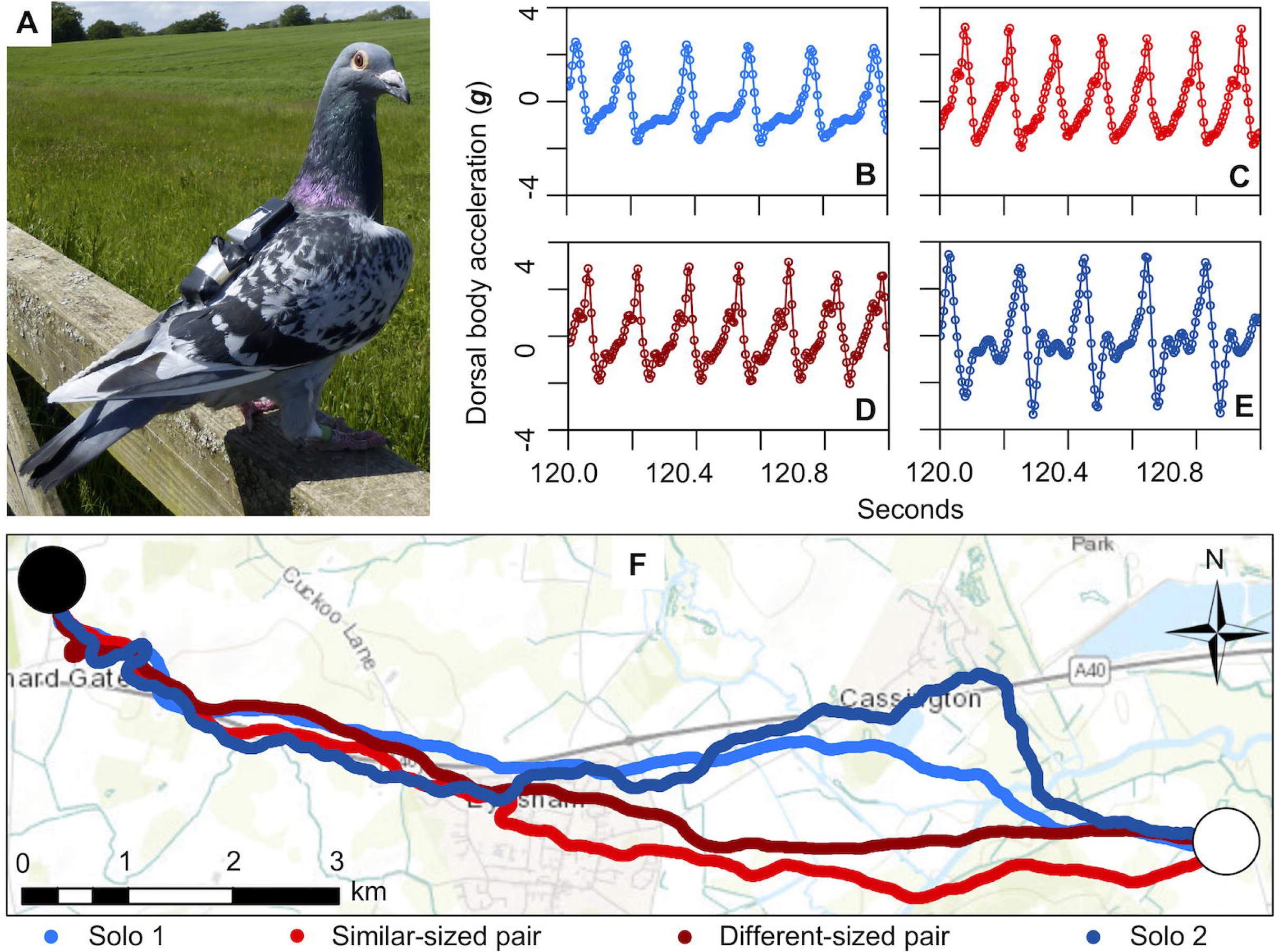
Examples of accelerometer and GPS data recorded during solo and paired flights. **A,** Bird S30 carrying an accelerometer (top) and GPS sensor (bottom) attached via Velcro strips. **B-E**, Dorsal body (DB) acceleration recorded by the accelerometer during S30’s final release in each of the four conditions: **B**, solo flight (blue); **C**, paired flight with a similarsized bird (red); **D**, paired flight with a different-sized bird (dark red) and **E**, solo flight (dark blue). Accelerometer data has been filtered and gravity removed (see Methods). Note the higher wingbeat frequency when the bird is flying in a pair. **F**, Routes flown by S30 during the final release of each of the four conditions (same flights as those shown in **B-E**). Note the straighter trajectory, and hence greater route accuracy, of the paired flights. Black circle corresponds to the release site and white circle corresponds to the home loft. Map designed using ArcGIS 10.4.1 (Esri Inc., Redlands, USA) using the World Topographic Map[23]. Scale bar shows 3 km.

**Figure 2.**
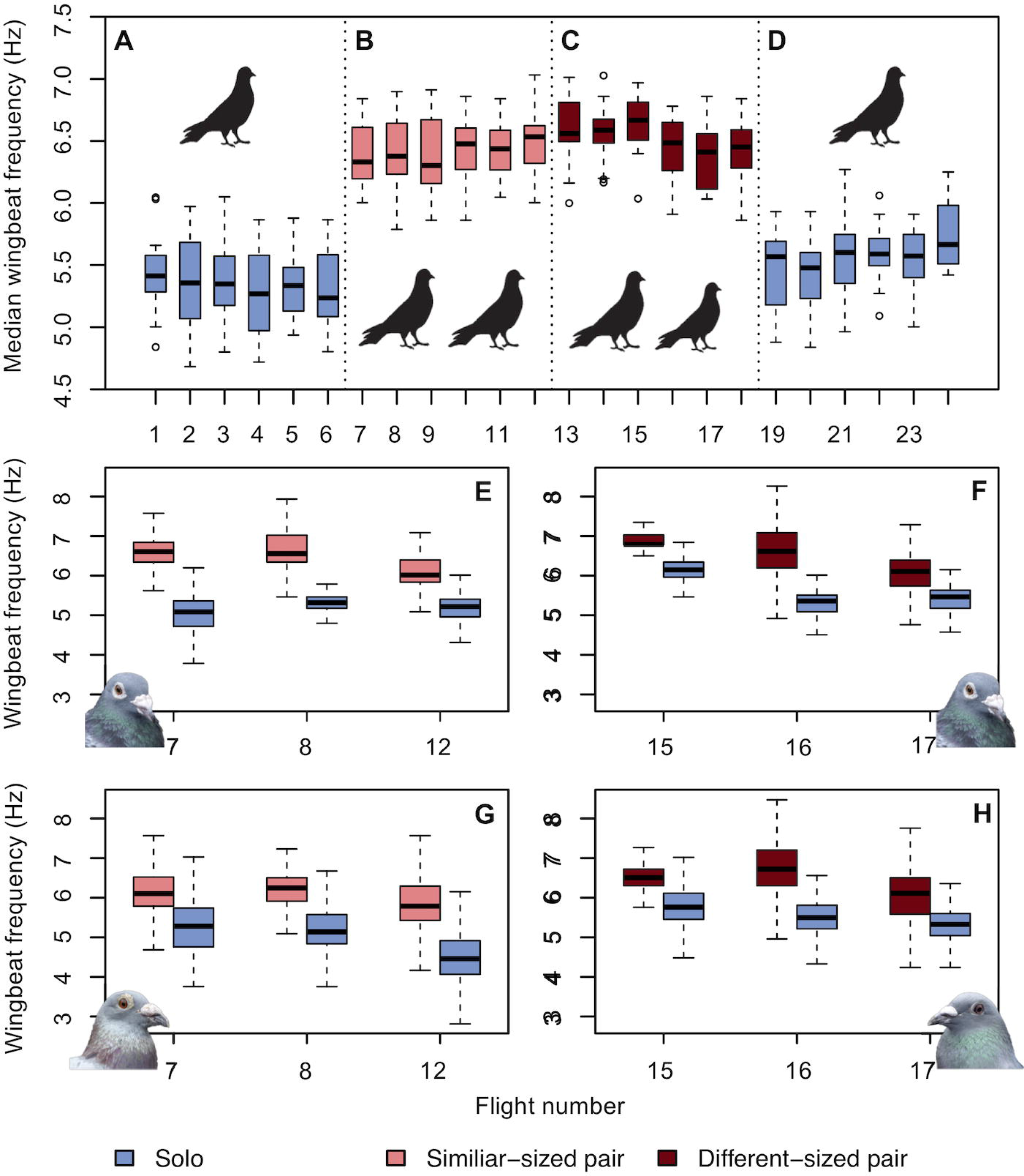
Wingbeat frequency as a function of flight number. **A-D**, Median wingbeat frequency (raw data with no covariates) for all 20 birds during each of the four experimental phases: **A**, six individual releases (solo); **B**, six releases with a similar-sized bird (similar-sized pair); **C**, six releases with a different-sized bird (different-sized pair); and **D**, six individual releases (solo). **E-H**, Raw wingbeat frequency data for birds which flew in a pair (< 50 m distance) and solo (> 300 m) during the same flight. **E** and **G** are from the similar sized pair of birds B01 and B82, respectively (pictured). **F** and **H** are from the different-sized pair B01 and B07 (pictured). Boxplots show the median and upper and lower quartiles, and whiskers correspond to 1.5 times the interquartile range.

## Results

We analysed the data from all flights using Bayesian hierarchical models to account for variation due to a set of environmental covariates, and the individual identity of the focal bird and any partner (see Methods). Our results show that when birds flew in pairs, their median wingbeat frequency increased by 1.00 Hz (95% Bayesian credible interval (CrI) [0.61, 1.38]) relative to flying solo, representing an increase of 18.2% (Fig. 2B-C). This was not associated with greater variability in wingbeat frequency, the standard deviation of which remained stable between solo and paired flight (2% lower standard deviation for size-matched pairs). Likewise, the median peak-to-peak amplitude of dorsal body (DB) acceleration was similar in both solo and paired flights (difference of 0.16 *g*, where *g* is gravitational acceleration; 95% CrI [-0.16, 0.47], where the fact that the credible interval crosses zero indicates that any difference is statistically indistinguishable from zero). This combination of increased wingbeat frequency and unchanged amplitude of DB acceleration resulted in a net 22.5% reduction in the median peak-to-peak amplitude of dorsal body (DB) displacement (-4.65 mm, 95% CrI [-6.31, -3.06]) through the wingbeat, as a result of the correspondingly shorter time period over which DB acceleration is integrated to produce DB displacement (see supplementary text for further analysis of the oscillatory accelerations experienced by an accelerometer).

These statistical findings were consistent within and between pairs, irrespective of whether the birds were flying in similar-sized or different-sized pairs (0.01 Hz increase per mm difference in tarsus length between the pair; 95% CrI [-0.13, 0.16]; Fig. S1), or whether the bird was in front or behind during the flight (0.01 Hz increase for travelling behind, 95% CrI [-0.10, 0.12]). Moreover, the probability of whether an individual bird flew ahead in a given pair was unaffected by the birds’ tarsus length (0.01 per mm difference in tarsus length, 95% CrI [-2.68, 2.44]), solo airspeed (-0.22 per m s^-1^ difference in median solo airspeed, 95% CrI [-3.08, 2.65]) or body mass (0.02 per g difference in body mass, 95% CrI [-0.10, 0.15]; Fig. S4), meaning there is no evidence that the larger or faster bird sets the pace by leading. Interestingly, and in contrast to the closely coordinated flight of birds flying in V-formation[12], there was no correspondence between the front and back bird’s median wingbeat frequency (β = -0.23, 95% CrI [-1.23, 0.75]; Fig. S3), indicating that their wingbeats cannot have been phase-locked through most of the flight. All of these results were computed after accounting for the birds’ median airspeed, date of release and weather variables, none of which had an effect on wingbeat frequency that was distinguishable from zero (Fig. S2, Table S1). Hence, whilst we found no evidence in support of our hypothesis that birds of different sizes would specifically have to adjust their wingbeat frequency or amplitude to stay together as a pair, we found clear evidence that both birds in a pair increased their wingbeat frequency, independent of individual size or solo flight speed.

In addition to these results for all releases, one similar-sized pair and one different-sized pair separated during three releases each, which means we can fortuitously compare sections of paired and solo flight within the same release. The results for these six releases clearly confirm that wingbeat frequency increases as a direct result of flying in a pair, because the birds’ median wingbeat frequency decreased by 1.01 ± 0.30 Hz (mean ± s.d.) after they separated and flew solo (raw values with no covariates; Fig. 2E-H).

As previous research has shown that pigeons increase their wingbeat frequency by up to 0.1 Hz as flock density increases [14], we analysed the effect of horizontal inter-individual distance ranging from 0 m (i.e. directly above or below another bird) to 50 m (i.e. the cut off point for flying in a pair) in a random sample of 100 wingbeats from each flight. This subsampling was necessary due to the computational demands of dealing with the otherwise extremely large volume of data. In total, we analysed 45,500 wingbeats from solo and paired flights. The results demonstrate that birds flying with no horizontal spacing did indeed have the highest wingbeat frequency (increase of 1.21 Hz relative to flying solo; 95% CrI [0.81, 1.61], 21.6%), with wingbeat frequency decreasing by 0.011 Hz for every metre increase in horizontal spacing (95% CrI [-0.012, -0.009]). Thus, birds flying 50 m apart had an expected wingbeat frequency 0.54 Hz lower than birds flying 0 m apart. Nevertheless, the act of flying in a pair still had a larger overall effect than the distance between (or density of) the birds, which meant that even birds flying 50 m apart increased their wingbeat frequency by 0.66 Hz (11.9%) relative to flying solo. On average, birds flew with a median spacing 12.12 ± 4.76 metres (mean of the means for all pairs ± standard deviation), which equates to a 1.07 Hz increase in wingbeat frequency under the fitted relationship (*cf.* the 1.00 Hz increase in median wingbeat frequency was over the whole flight).

To explore the mechanism underlying this change in wingbeat frequency, we divided each wingbeat into an upstroke and a downstroke phase. We defined these phases with respect to the peaks and troughs of the DB acceleration, which results from a combination of aerodynamic and inertial forcing (see supplementary text for further detail). Whereas the dorsal aerodynamic force is expected to peak mid-downstroke when the wing reaches its maximum flapping speed, the dorsal inertial force is expected to peak at the start of the downstroke when the wing’s downwards acceleration is maximal. It follows that the maximum DB acceleration will be reached somewhere between the start and middle of the kinematic downstroke, and similarly for the minimum, which will be reached somewhere between the start and middle of the kinematic upstroke. Hence, the downstroke phase, which we define as running from the point of maximum to minimum DB acceleration, is expected to lag the kinematic downstroke slightly (and similarly for the upstroke), but by less than a quarter of a cycle. With these definitions, we found that birds reduced the median duration of the upstroke phase by 20.6% (-27.99 milliseconds, 95% CrI [-35.61, -19.92]) when flying in pairs, whereas the median duration of the downstroke phase did not vary significantly (-3.53 milliseconds, 95% CrI [-8.35, 1.38]; 2c-d). It is clear by inspection of the wingbeat acceleration traces that this decrease in upstroke duration results in a less asymmetric pattern of forces between the two wingbeat phases (compare red versus blue lines in Fig. 1B-E), so that this change in wingbeat frequency essentially represents a switching of kinematic - if not aerodynamic - gait [24].

We hypothesise that a potential function of increasing wingbeat frequency and decreasing oscillatory displacement of the body may be to enhance visual stability when attending to nearby conspecifics. We therefore conducted a second experiment using head-mounted accelerometers on six homing pigeons on short-range flights (950 m), flying solo and in pairs, to determine if the same measured changes in wingbeat characteristics result in increased head stability (see Methods). In close agreement with the first experiment, birds flying in pairs increased their median wingbeat frequency by a mean of 1.10 Hz ± 0.26 relative to flying solo (6.6 ± 0.42 Hz mean ± s.d. for pairs; 5.5 ± 0.46 Hz for solo). More importantly, however, the results also show that the median peak-to-peak head displacement simultaneously decreased by 5.3 ×10^-3^ m ± 6.6 × 10^-4^ between solo and paired flight, representing a 30% reduction in the amplitude of oscillatory head displacement (Fig. 3). This substantial improvement in translational head stability is expected to result in a significant reduction in the retinal slip of nearby objects including flight partners.

**Figure 3.**
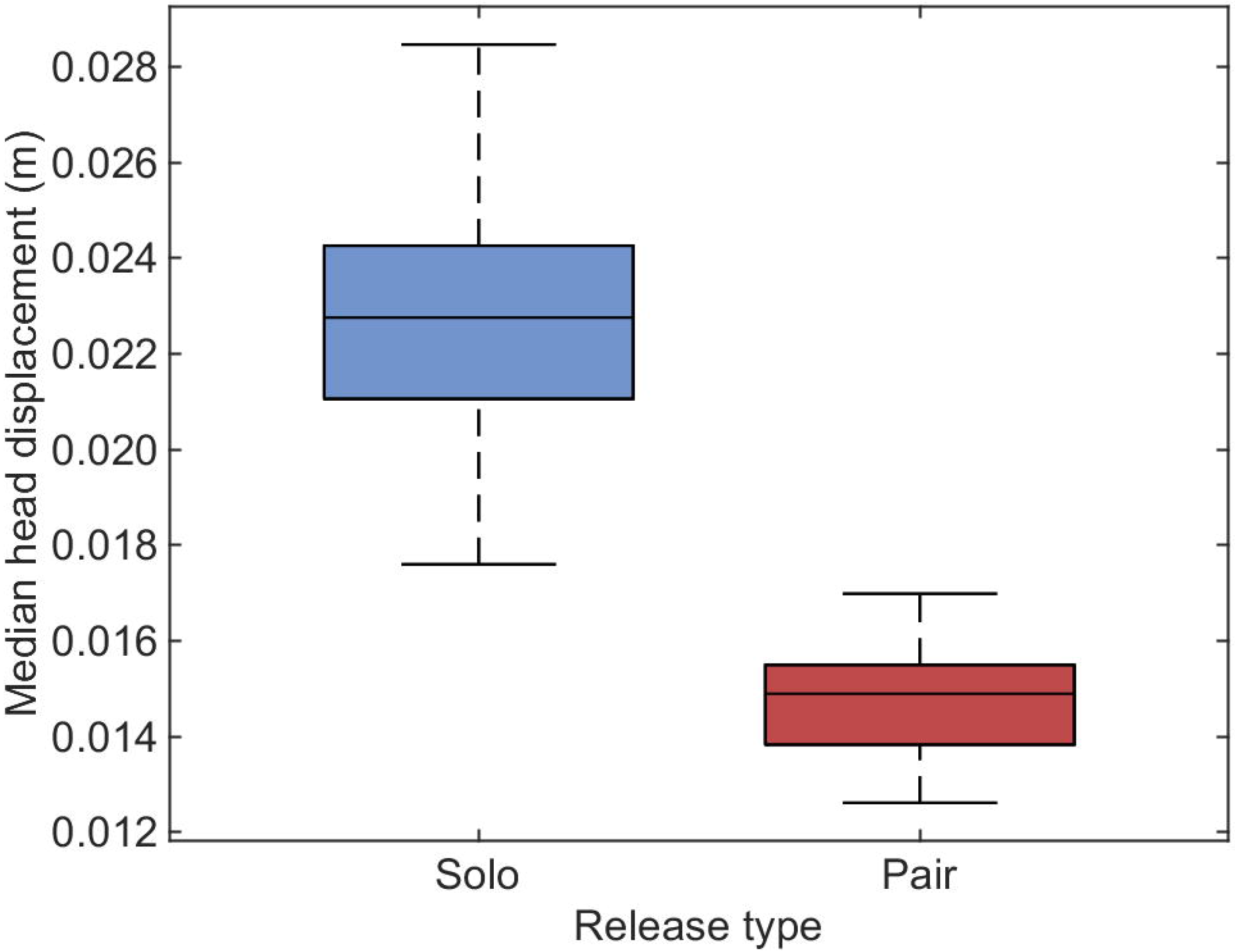
Boxplot of median vertical peak-to-peak head displacement across six birds, each flying once solo and once in a pair. Bottom and top edges of the box indicate the 25th and 75th percentiles and the whiskers extend to the most extreme data points. Note that there is no overlap between the median head displacements in the two conditions.

In summary, by reducing the duration of their upstroke phase, birds flying in a pair were able to accommodate one additional wingbeat per second, whilst maintaining the same peak-to-peak DB acceleration and simultaneously increasing the vertical stability of the head. Intuitively, a higher-frequency kinematic gait adopted in paired flight will therefore be associated with a higher mechanical power input than the lower-frequency flight kinematic gait adopted in solo flight. Of course, a higher mechanical power requirement in paired flight could still be associated with a lower cost of transport if this increased frequency were more than compensated by an increased flight speed. However, whereas birds migrating in V-formations are known to increase their airspeed as flock size increases [25], the birds in our study only increased their airspeed by 3.3% when flying in pairs (0.64 m s^-1^ increase, 95% CrI [0.08, 1.2]). As we now explain, this increase in airspeed is much smaller than could have been expected to be caused by the increase in wingbeat frequency alone, suggesting that there must have been other compensatory changes in the kinematics.

In cruising flight, the net thrust of the wings balances the drag on the body, which scales as *ρU*^2^*S*_*b*_ where *ρ* is air density and *S*_*b*_ is body frontal area. For a 3% increase in airspeed *U*, it follows that the time-averaged thrust can only have increased by just over 6%. In contrast, the thrust on a flapping wing has been shown to scale as *T*~*ρS*_*w*_*f*^2^*A*^2^, where *S*_*w*_ is wing area, *f* is wingbeat frequency, and *A* is wingbeat amplitude [26]. Assuming all other things are equal, an 18% increase in wingbeat frequency would therefore be expected to produce about a 39% increase in thrust. However, we also know that the time-averaged lift must balance the bird’s weight when cruising, and that lift scales as *L*~*ρS*_*w*_*U*^2^ in fast forward flight when the contribution of the wing’s own flapping speed can be ignored. This implies that the 3% increase in airspeed (i.e 6% increase in *U*^2^) would have to have been countered by either a 6% decrease in wing area, or an equivalent decrease in the proportionality constant of the scaling relationship (i.e. the wing lift coefficient). Either kinematic change would be expected to attenuate the thrust similarly, thereby reducing its expected increase to approximately 31%. This is still significantly higher than the 6% increase in thrust estimated on the basis of the 3% increase in airspeed, but the scaling *T*~*ρS*_*w*_*f*^2^*A*^2^ implies that these figures could be brought into line by an accompanying 10% decrease in wingbeat amplitude (see supplementary text for further confirmation).

The aerodynamic power requirement of a flapping wing has been shown to scale as *P*_*A*_~*ρcS*_*w*_*f*^3^*A*^2^ where *c* is the wing chord [26]. The inertial power requirement also varies in proportion to *f*^3^*A*^2^, albeit with some further complications related to the effect of varying wing span, but is an order of magnitude smaller than the aerodynamic power requirement [27,28] so is neglected here for simplicity. Assuming that the 18% increase in wingbeat frequency *f* between solo and paired flight was accompanied by a 6% decrease in wing area *S*_*w*_ and by a 10% decrease in wingbeat amplitude *A* as required to meet the equilibrium conditions above, then the aerodynamic power requirement (in J s^-1^) would have increased by approximately 25% when flying in a pair. Given the 3% increase in airspeed, it follows that the aerodynamic cost of transport (in J m^-1^) must also have increased by some 21%. However, another key benefit often ascribed to flying in flocks is the ability to pool navigational knowledge. This should improve homing route accuracy [7,8], which could offset the increased aerodynamic cost of transport and increased aerodynamic power requirement by simultaneously decreasing the distance and duration of the flight.

We calculated the birds’ route accuracy flying solo and in pairs using a weighted mean cosine of the angle between the birds’ heading and destination. Flying in a pair resulted in a 6.9% increase in route accuracy relative to both the Phase 1 and Phase 4 solo releases (0.06, 95% CrI [0.01, 0.10]; Fig. 1), and a 6.5% decrease in route length (Table S2). This is consistent both with theory[7,8] and with previous empirical studies [5,6]. Offsetting the 21% higher cost of transport when flying in a pair against the 6.5% reduction in route length, we would expect a net increase of approximately 14% in the total mechanical energy expended when flying home to the loft in a pair. The increase in the total metabolic energy expended could be higher or lower than this, depending upon whether and how the efficiency of the flight muscles varies with flapping frequency, but it is reasonable to assume that an increase in mechanical energy would also be associated with an increase in metabolic energy.

Whilst these figures are necessarily approximate, the qualitative conclusion of this analysis - that the energy expended returning to the loft would have been higher in paired than in solo flight - is robust to the uncertainty in our estimates of the changes in wingbeat frequency and airspeed. Substituting the limits of the 95% CrI’s for these variables into the preceding calculations leads to an estimated 4% or 24% increase in the total energy expended in paired flight if the variables fall at their respective upper or lower limits, and an estimated 10% or 17% increase if they fall at opposite ends of their limits. These are worst-case figures for how the uncertainties could combine, all of which lead to the conclusion that flying in pairs is associated with a substantial net increase in energy consumption over the flight. As we now discuss, this surprising result indicates: that (*i*) the act of flying in a pair necessitates birds to alter their wing kinematics to a higher-frequency kinematic gait; and that either (*ii*) the navigational benefits of flying in a pair are sufficient to outweigh the increased cost of transport over longer homing distances, such that flying in pairs makes sense as a general homing strategy, or (*iii*) there are other benefits of paired flight that outweigh its net energetic cost even over short distances.

## Discussion

One of the most commonly cited reasons for travelling as a group is to reduce energy expenditure and enhance locomotor performance [11,12]. Previous research in cluster flocking pigeons has shown that the energetic cost of flocking increases slightly with increasing flock density [14]. However, this study did not compare the cost of flying in a cluster flock relative to the alternative of flying solo, which means, as we now show, that previous work has inadvertently understated the energetic costs of cluster flocking by an order of magnitude. Specifically, whereas pigeons have previously been shown to increase their wingbeat frequency by a mere 0.1 Hz with increasing flock density [14], our results show that the very act of flying with another bird increases a pigeon's wingbeat frequency by 1.0 Hz (18%), which results in an estimated 21% increase in the aerodynamic cost of transport. Although birds flying in pairs were simultaneously able to offset some of the energetic cost by flying more accurate routes home, the increases in route accuracy and airspeed were insufficient to compensate for the increased aerodynamic power requirements, which resulted in a net energetic loss on the order of 14% when flying moderate distances (~7 km) together in a cluster formation. Moreover, the fact that pigeons flying in pairs display a 18% increase in wingbeat frequency over solo flight suggests that the majority of the additional cost comes merely from the act of flying with another individual, rather than from the density of the flock, the relative spatial position of the bird, or the size of its partner. Indeed, the size of a bird’s partner, and whether that bird was a leader or follower, had almost no effect on its measured wingbeat pattern. Even so, differences in inter-individual horizontal spacing did result in a 0.54 Hz difference in wingbeat frequency between birds travelling 0 to 50 m apart, with this increase ranging from 11.9 to 21.6 %, respectively. Thus, the act of flying with a conspecific resulted in a substantial alteration of the wingbeat - even the adoption of a different kinematic gait. As we now explain, not only does this earlier omission mean that the costs of flocking have been massively underestimated - it also means that their mechanistic origin must be re-evaluated.

Two key hypotheses have been proposed for the increase in wingbeat frequency seen in denser cluster flocks: (*i*) negative aerodynamic interactions between flock members and (*ii*) increased need for control and collision avoidance [14]. Whereas a focus on the small effects of spacing within a flock led previous work to hypothesise a possible aerodynamic basis to the costs of cluster-flocking [14], our work clearly demonstrates that both birds within a pair increase their wingbeat frequency, which suggests that these effects are unlikely to have been related to negative aerodynamic flow interactions, since the bird in front does not fly in the wake of the bird behind. On the other hand, higher wingbeat frequencies can be used to enhance both stability and manoeuvrability [14,29–31]. We therefore hypothesise that the increase in wingbeat frequency is related to paired flight necessitating a greater degree of control, which could come about in two different ways. First, flying with conspecifics may require enhanced manoeuvrability and control because birds need to adjust their orientation continuously and rapidly, both to stay together and to avoid collisions [14]. Second, birds may require enhanced visual stability when flocking, in order to observe and coordinate with individuals whose proximity makes the effects of motion parallax significant [32,33].

Birds make kinematic control inputs on a wingbeat-to-wingbeat basis, so increasing wingbeat frequency will increase the rate at which control inputs can be made, enhancing the bird’s ability to respond to the movements of others and increasing the precision of its response. Moreover, increased wingbeat frequency is expected to amplify flight stability [31], which could both enhance control and visual stability. Unlike humans, birds have a limited range of eye movement and therefore visual stabilisation is facilitated by compensatory motion of the sophisticated avian head-neck system and is mediated by visual, vestibular and proprioceptive cues [34,35]. Without image stabilisation mechanisms, birds would have difficulty differentiating the motion of a target or obstacle from head or body motions, which is especially problematic when viewing nearby objects or conspecifics. Our results show that dorsal body displacement in reaction to the wingbeat is attenuated by 23% in the higher-frequency kinematic gait adopted in paired flight (see supplementary text for theoretical analysis), which should naturally translate into a reduced amplitude of head motion. Indeed, using data from head-mounted accelerometers, we prove that the heads of birds flying in pairs experience significantly less vertical head displacement relative to flying solo (~30%). Pigeons flying in pairs have also been recorded to reduce their angular head saccades relative to flying solo, which suggests either an increased focus on their partner or a decreased focus on the environment [36]. Furthermore, the linearly declining effect of paired flight with increasing inter-individual distance suggests that the observed changes in wingbeat frequency are less to do with the local aerodynamic environment, the effects of which drop off sharply, and are instead related to the coordination of flock flight, noting that the effects of motion parallax decline linearly with distance. Together, our results suggest that a higher wingbeat frequency may be prerequisite for flocking flight because of the increased demands for stability and control.

Managing energy expenditure is critical for survival, and is a primary focus for natural selection. While our results demonstrate that birds can derive navigational benefits from flying in pairs even during short-range flights along familiar routes, the 7% increase in route accuracy over this range was apparently insufficient to counterbalance the cost of the 18% increase in wingbeat frequency, with an estimated increase in mechanical energy expenditure on the order of 14%. Despite this, only six releases had to be repeated due to birds separating at the start, and only twelve out of 116 pair releases (10%) resulted in separation part-way through the release. Hence, the observed preference for paired flight suggests that either (*i*) the general strategy of flocking is adaptive because the navigational benefits of flocking are sufficient to outweigh the increased cost of transport when homing over longer distances, or (*ii*) the other benefits of flocking, such as predator protection, outweigh the increase in energy expenditure required to fly in pairs, even over quite short distances.

Although minimizing energy consumption may not be an especially strong selection pressure for homing pigeons that have been selectively bred to return quickly to the loft, and which have *ad libitum* access to feed, the results of our study nevertheless indicate that flying with conspecifics entails an energetically expensive alteration to wingbeat kinematics. As many other species of birds also preferentially fly in cluster flocks, our results suggest that the additional benefits of flocking must outweigh any accompanying increase in energy expenditure. The overall 9% reduction in homing flight time that we observed represents a 9% reduction in the period over which our birds were exposed to predation risk when returning to the loft. Moreover, not only does the act of flying in a pair dilute the chance of fatality during a predation event by 50%, but the probability that such a predation event is successful decreases as flock size increases, presumably through a combination of increased opportunity for vigilance and predator confusion effects [37]. Therefore, for pigeons, the ultimate benefits of flocking, such as protection from predators and the pooling of navigational knowledge, must together outweigh the energetic cost of flying with conspecifics.

Over longer flight distances or circumstances where an individual has substantially less navigational knowledge than the flock, it is possible that the navigational benefits of flocking might be sufficient to produce a net reduction in the amount of energy expended despite the increased aerodynamic power requirement that we report. In order to gain energetic savings from a 21% increase in the cost of transport (J m^-1^), the homing distance (m) would have to be >17% shorter when flying in a pair to result in net energetic savings (J). In this scenario, birds would experience an increased rate of energy expenditure which would be compensated by net energetic savings and a reduced risk of predation. Either way, the birds in our study still opted to fly in pairs despite collective travel resulting in an energetic loss at the individual level.

Overall, the results of our study of a cluster flocking species stand in contrast to previous studies of birds flying in V-formations [11–13]. Formation flight is typically utilised by medium to large sized birds during goal-orientated movement, whereas cluster flock formations are utilised by smaller birds, such as starlings, in movement ranging from orientated to highly tortuous motion [15,16]. Bird size and the complexity of movement paths may both contribute to the observed differences in wingbeat patterns between flock formation types. Whilst birds flying in V-formations are able to fly in aerodynamically optimal positions to conserve energy, the naturally higher wingbeat frequencies of smaller birds, their smaller turning angles, and the rapidity of their directional changes may preclude flying in energy-saving formations and instead necessitate a wingbeat pattern that facilitates a greater degree of control. Thus, the demands of moving in irregular three-dimensional flocks may alter the way in which a bird flies. Overall, our results provide key new insights into both the biomechanical consequences of close cluster flocking, and the energetic investments that pigeons make to gain access to collective navigational knowledge and predator protection. Taken together, our results demonstrate that flocking is, for pigeons, both fundamentally important and fundamentally expensive.

## Materials and Methods

### a) Experiment 1

#### i) Subjects

Twenty homing pigeons aged 1 or 3 years were used. Body size was quantified by measuring tarsus length (mm) and body mass (g). Tarsus length was measured with callipers sensitive to 0.1 mm using the methods described in Sutherland et al. [22]. Body mass was measured using digital scales (Salter ARC Electronic Kitchen Scales, Salter, UK; ± 1 g). All subjects completed a minimum of 15 solo flights from the release site used in this study immediately preceding the start of the experiment. The subjects were housed with ~120 other pigeons in two neighbouring lofts at the Oxford University Field Station, Wytham, UK (51°46’58.2”N, 1°19’2.7”W). Access to water, grit and a standard pigeon feed mix were available ad libitum at all times in the loft. The protocols outlined in this paper were approved by the Local Ethical Review Committee of the University of Oxford’s Department of Zoology.

#### ii) Data logging

The birds were tracked using 5 Hz GPS loggers (QStarz BT-Q1300ST, 15 g) and 200 Hz tri-axial accelerometers (± 16 *g*; Axivity AX3, 11 g), which were attached via Velcro strips glued to trimmed feathers on the birds’ back. In total, the loggers and fastenings weighed 27 g. To enable subjects to adapt to carrying the additional mass, clay weights were attached to them throughout the pre-training and experimental periods, which meant the weights were attached for a minimum of 43 days prior to the start of the experiment. The weights were exchanged for the loggers immediately prior to each release. GPS and accelerometer data were downloaded using QTravel (Qstarz International Co., Ltd., Taipei, Taiwan; version 1.48(T)) and Open Movement (Om) GUI Application (Newcastle University, UK; version 1.0.0.28), respectively.

The weather, including mean wind speed per minute (ms^-1^), a running mean of the wind bearing over the previous ten minutes, temperature (°C), humidity (%) and barometric pressure (hPa), were recorded using a WS2083 Professional Wireless Weather Station with USB upload (Aercus Instruments, UK) situated 5.5 m above the ground near the pigeon lofts and Cumulus Weather Station Software (Sandaysoft; version 1.9.4).

#### iii) Experimental procedures

The release site was located 7.06 km from the loft on a bearing of 282° (Barnard Gate; 51°47’48.1”N, 1°25’3.3”W). The experiment consisted of four phases: Phase 1 - six individual releases (solo 1); Phase 2 - six releases with a bird of a similar size (similar-sized pair); Phase 3- six releases with a bird of a different size (different-sized pair); and Phase 4 - six individual releases (solo 2). Bird pairings can be found in Table S3. Releases were conducted between June and September 2015, on days when the sun was visible and the wind speed was < 7 ms^-1^. Subjects participated in a maximum of two releases per day, with a minimum of three hours between each release. The birds had to complete a minimum of one third of the flight together for the flight to be included in the analysis. If the birds spent less than one third of the flight together, the flight was repeated. In total, six releases out of 116 pair releases had to be repeated. One different-sized pair did not complete the final pair release after repeatedly separating. The Velcro failed on bird S27 after the third release with a different-sized bird therefore the pairing S27 and S84 only completed three different size pair releases and S27 did not complete the final solos. In addition, S13 only completed one final solo before the Velcro failed, S87 completed four final solos, and S05 and S25 completed five final solos each. The remaining 15 birds all completed the final solo releases.

#### iv) Data processing

Data were processed using the procedures outlined in Taylor *et al.* [20]. For each GPS point, the orthodromic (great-circular) distance travelled and birds’ final bearing from the previous point were calculated using the haversine formula and forward azimuth, respectively. The dorsal accelerometer measurements were filtered by taking a running mean over three data points (0.015 s). Static acceleration (or gravity) was removed by subtracting a running mean over 15 wingbeat cycles (> 2 s; Fig S5). The wingbeat frequency (number of wingbeats per second; Hz) and peak-to-peak dorsal body (DB) acceleration (*g*) using the dorsal acceleration signal (Z-axis) were calculated for each individual wingbeat. The amplitude of the DB displacement (mm) was then calculated by the double integration of dorsal accelerometer measurements [14,20]. In addition, we calculated the duration of the “downstroke” from the peak downstroke force (maximum *g*-force) to the lower reversal point (minimum *g*-force). The “upstroke phase” duration, which included the start of the downstroke, was measured from minimum *g*-force to the maximum. We used the maximum and minimum *g*-force peaks to divide the wingbeat for consistency, as the start of the kinematic downstroke was not distinguishable in the data from paired flights. See supplementary text for further analysis.

Wind support, crosswind and airspeed were calculated using the methods described in Safi et al.[38] using the measurements from the weather station and speed derived from the GPS devices. Humid air density (kg m^-3^; *ρ*_*air*_) was calculated from measures of barometric pressure (hPa; *P*), temperature (°C; *T*_*c*_) and relative humidity (%; ϕ) recorded by the weather station, using the following calculation derived from the ideal gas law:

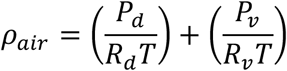

where *P*_*d*_ is pressure of dry air (Pa), *R*_*d*_ is gas constant for dry air [287.05 J/(kg * K)], *P*_*v*_ is pressure of water vapour (Pa), *R*_*v*_ is gas constant for water vapour [461.495J/(kg * K)] and *T* is ambient temperature (K). *P*_*v*_ can be calculated from the saturation of vapour pressure (*P*_*sat*_) and relative humidity (ϕ):

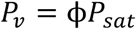

We used the Arden-Buck [39,40] equation to calculate *P*_*sat*_, where *P*_*sat*_ (hPa) is calculated as:

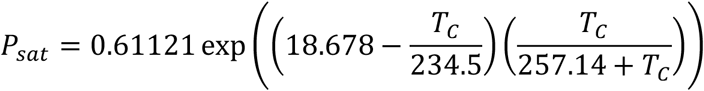

*P*_*d*_ can then be calculated from the barometric pressure (*P*) and the vapour pressure of water (*P*_*v*_):

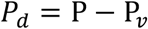

The birds’ route accuracy was calculated using a weighted mean cosine of the angle (*θ*) between the birds bearing and the bearing to the loft for each timestep, where *θ* is equal to the smallest angle difference so that *θ* ranged from 0 (heading directly to the loft) and 180 (heading directly away from the loft), and the orthodromic distance between each GPS point (*d*) using the following calculation:

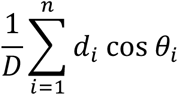

where *D* is the total distance flown. Route accuracy is, therefore, on a scale of -1 (heading in a straight-line away from the destination) to 1 (straight-line to the destination). For orientated movement, route accuracy is >0, which means route accuracy is akin to the straight-line index [41], but enables us to calculate the accuracy for sections of a flight, rather than a whole flight, which is necessary if the birds separate during the flight and fly solo.

The data were trimmed within a 200m radius around the release site and the loft to remove take-off and landing. When analysing the pair tracks, sections of flight where the birds were ≥ 50 m apart was excluded. In addition, if the birds swapped front-vs-back positions, the bird who spent the majority of the flight in front based on GPS positioning was identified and the rest of the data from when the other bird was in front was excluded.

#### v) Data analysis

We analysed the data using Bayesian hierarchical models, which are analogous to mixed models in frequentist methods and enabled us to account for the effects of each bird both as an individual and a partner in a specific pair. The median wingbeat frequency 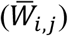 for the pair (*i,j*) was assumed to be normally distributed,

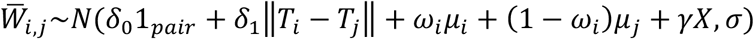

where 1_*pair*_ is an indicator variable equal to 1 if the bird flew in a pair, and 0 for solo flights. *δ*_0_ is the difference in wingbeat frequency between the solo and paired flight. *δ*_1_ is the difference in wingbeat frequency for every mm absolute difference in tarsus length (*T*) between the pair (*i,j*). For solo flights, the term involving *δ* equals zero. The expression *ω*_i_*μ*_*i*_ + (1 - *ω*_*i*_)*μ*_*j*_ represents a weighted average of the solo wingbeat frequency (*μ*) of birds *i* and *j* with a mixing weighting (*ω*_i_), which determines the weight placed on the bird’s own solo wingbeat frequency (*μ*_*i*_) relative to that of its partner (*μ*_*j*_). For solo flights, this weighted average equals bird *i*’s solo wingbeat frequency (*μ*_*i*_). The weighting is bounded to lie between 0 and 1 and was determined by a logistic sigmoid function of the absolute difference in tarsus length,

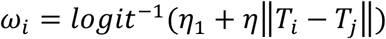

However, across all cases, there was no consistent effect of tarsus difference on the mixing weighting.

Finally, *γX* represents the effect of the covariates, which accounts for median wind support (m s^-1^), median crosswind (m s^-1^), median temperature (°C), median humidity (%), humid air density (kg m^-3^) and the date of release treated as a categorical variable. The birds’ median airspeed (m s^-1^) was also added as a covariate on all models except for models of airspeed. We used airspeed rather than ground speed as a covariate because ground speed and wind support were correlated (Fig. S5). In terms of the response variable, there was almost no difference between the models of airspeed and ground speed as the model accounts for the effect of wind (0.64, 95% CrI [0.08, 1.20] compared to 0.69, 95% CrI [0.10, 1.28]). For consistency with the covariates, we present the results of the model for airspeed. A covariate indicating whether the bird was a leader or follower was also added to the model in a secondary analysis to determine the effect of the birds’ position on wingbeat frequency.

In addition to modelling the median values, we also took a random sample of 100 individual wingbeats to analyse the effect of horizontal distance between birds in pairs. We analysed horizontal distance rather than three-dimensional distance because GPS precision is generally poorer in the vertical than the horizontal [42]. Horizontal distance (m) was added as a covariate to the paired data, along with a categorical covariate identifying the specific bird and flight to account for the repeated measures of 50 wingbeats from one flight. In total, 44,500 wingbeats from 454 unique bird and flight combinations were analysed.

To investigate whether the birds were flying in phase, we used the following model to identify whether the median wingbeat frequency of the follower 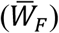 is related to the leader 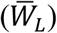 in pair (*P*):

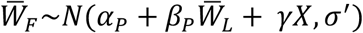

We also investigated whether the difference between the leader and follower’s tarsus length, body mass or solo airspeed (*S*) predicted why the bird was a leader (*L*) in the pair using a Bernoulli regression:

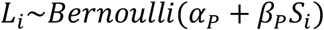

The model priors were centred on the null hypothesis using the mean, standard deviation and square root standard deviation of the solo data (Table S4). Eight Markov chain Monte Carlo (MCMC) chains were run simultaneously, each with 12,500 warm-up and 12,500 model iterations, which resulted in 100,000 samples for each posterior distribution. For the model involving raw wingbeat data, the model was run for 10,000 samples due to the size of the model. Across all estimated models and parameters, we detected convergence in the sampling distribution as determined by using a criterion 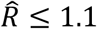 on all parameters[43]. The number of divergent iterations was 0.0-1.3% of the total sample size. The code for the wingbeat frequency model and the model output can be found in the Supplementary Materials.

In addition to these results, one similar-sized pair (birds B01 and B82 with a tarsus length of 32.4 mm and 33.3 mm, respectively) and one different-sized pair (birds B01 and B07 with a tarsus length of 32.4 mm and 35.2 mm, respectively) split and flew solo for more than 30% of the flight for three of their six releases. As the sample size is low, only descriptive statistics can be performed comparing the paired (< 50 m distance between birds) and solo flight (> 300 m distance).

We approximated the number of wingbeats difference between the solo and paired flight using the total flight distance (excluding the 200m take-off and landing) and the model results from route accuracy, airspeed and wingbeat frequency (Table S2). We calculated the number of wingbeats difference as not all of birds completed 100% of the paired flights together or with one leader.

Data processing and analysis were conducted using MATLAB (MathWorks, Natick, USA; version R2015a) and the open-source software R (version 3.4.2) [44]. Bayesian models were written in Stan [45] using the R interface RStan (version 2.16.2) [46].

### b) Experiment 2

#### i) Subjects

Six homing pigeons aged 3 to 10 years old were used. The pigeons were held under the same conditions as outlined above and all had experience of experimental releases and the release site. The protocols outlined in this paper were approved by the University of Oxford’s Zoology Animal Welfare Ethical Review Board (No. APA/1/5/ZOO/NASPA/Biro/ PigeonsHeadmountedsensors).

#### ii) Data logging

We used a custom-built ‘p-Sensor’ to simultaneously record head movement and position. The p-Sensor included an IMU with a combination of a tri-axial gyroscope, tri-axial accelerometer and tri-axial magnetometer recording at 60 Hz, and a GPS logger recording at 10 Hz. The IMU was mounted using double-sided tape onto a custom-made and custom-fitted wire mask designed to fit each bird’s head. The GPS logger, SD card, battery and microcomputer were placed in an elasticated backpack on the birds back. The instrumentation, mask, and backpack weighed 28.1 g and constituted 4.9 % of the body mass of the smallest bird, of which the IMU unit on the bird’s head only weighed 1 g. For more details, see Kano et al. [36].

All birds were habituated to wearing the custom-made mask for at least seven days prior to the flight. For each day of habituation, the bird was fitted with a mask and carefully monitored for two hours within its home loft for signs of discomfort and abnormal patterns of locomotion. After seven days of habitation in the loft, the pigeons were released outside the loft and allowed to fly freely under close observation.

#### iii) Experimental procedures

The release site was located 0.95 km from the loft on a bearing of 199° (Wytham Woods; 51°46’29.4”N, 1°19’18.7”W). The experiments were conducted on the 23^rd^ July 2017. Releases were only conducted when the wind was low (<5 m s^-1^) and the sun’s disc was visible. For the day of testing, the birds were fitted with a mask and allowed to habituate to wearing the mask in the home loft before being transported to the release site by car. The birds were released once solo and once in a pair on the same day. The release order was randomised.

#### iv) Data processing and analysis

The data processing was conducted as outlined in Taylor et al. [20]. Vertical (Z-axis) accelerometer measurements were smoothed by taking a running mean over five datapoints (0.083 s) and then filtered using a 4^th^ order high-pass Butterworth filter with a cut-off frequency of 1 Hz. The peak-to-peak vertical head displacement was calculated by the double integration of the vertical accelerometer measurements. We compared the median peak-to-peak vertical head displacement between solo and paired releases for each bird.

The raw data will be available on data dryad after acceptance.

## Supporting information

## Acknowledgments

We thank Lucy Larkman, Dave Wilson and Phil Smith for animal husbandry and technical support and Fumihiro Kano for his contributions to the development of the head-mounted sensor and mask for head attachment. We thank Marco Klein Heerenbrink, Oliver Padget, Jay Willis and Simon Taylor for helpful discussions. We thank Eoin Malins from the University of Oxford’s Doctoral Training Centre for providing computer access. L.A.T. and J.A.W. were funded by the Biotechnology and Biological Sciences Research Council (BBSRC) UK (grant number BB/J014427/1). B.L. was funded by Engineering and Physical Sciences Research Council (EPSRC) UK (grant number EP/F500394/1).

