## Supplementary material for "Birds invest wingbeats to keep a steady head and reap the ultimate benefits of flocking"

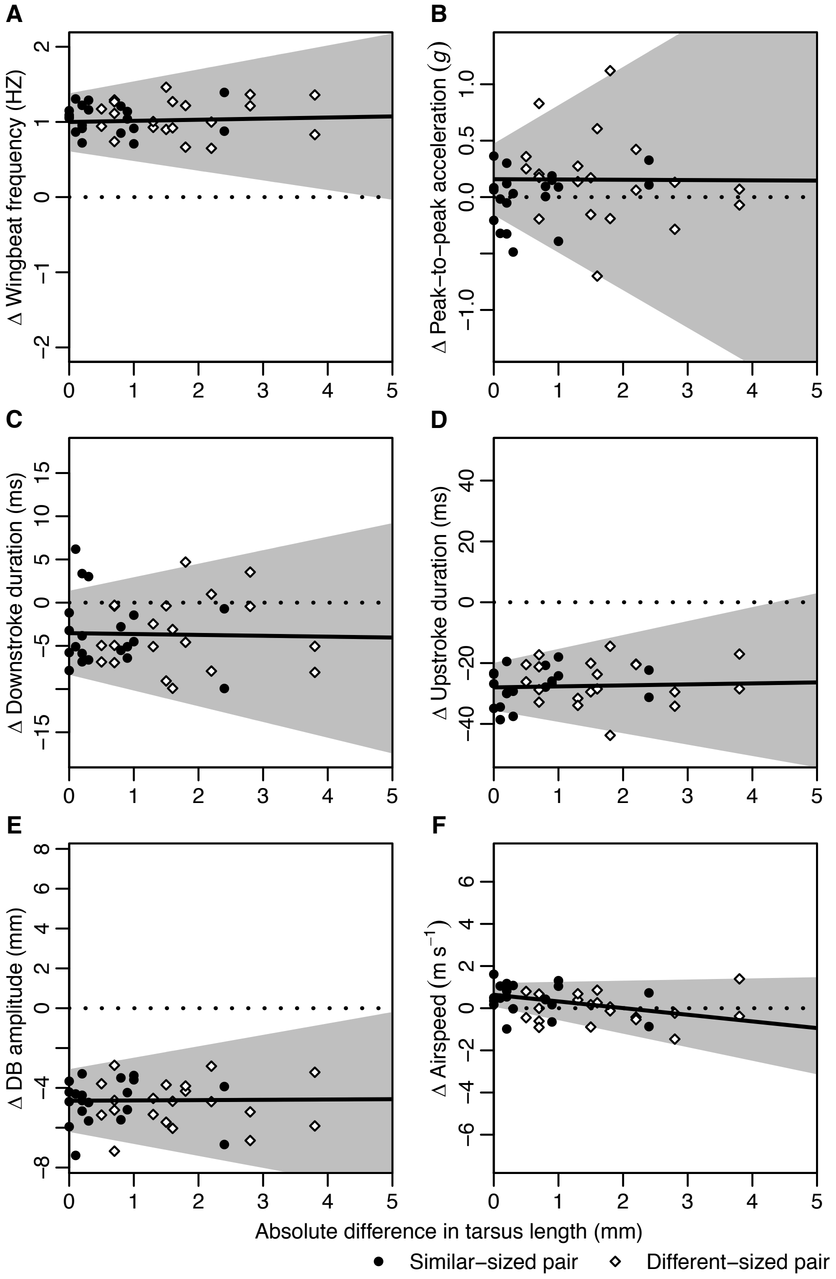


**Figure S1.** Bayesian hierarchical model results. Graphs show the difference (Δ) in (**A**) wingbeat frequency, (**B**) peak-to-peak dorsal body acceleration, (**C**) downstroke phase duration, (**D**) upstroke phase duration, (**E**) dorsal body (DB) amplitude and (**F**) airspeed between solo and paired flight per mm difference in tarsus length. Black line corresponds to the mean model fit which represents the difference between solo and paired flight per mm difference in tarsus length. Grey shaded area corresponds to the 95% credible interval of both the difference between solo and paired flight and the difference per mm difference in tarsus length. Points correspond to the difference between the median value for the paired flight and the median value for the solo flight, after accounting for the effects of weather, day and, in case of the a-c airspeed. Dotted line shows the solo value. All axes scaled to show ± 40% of mean solo value.

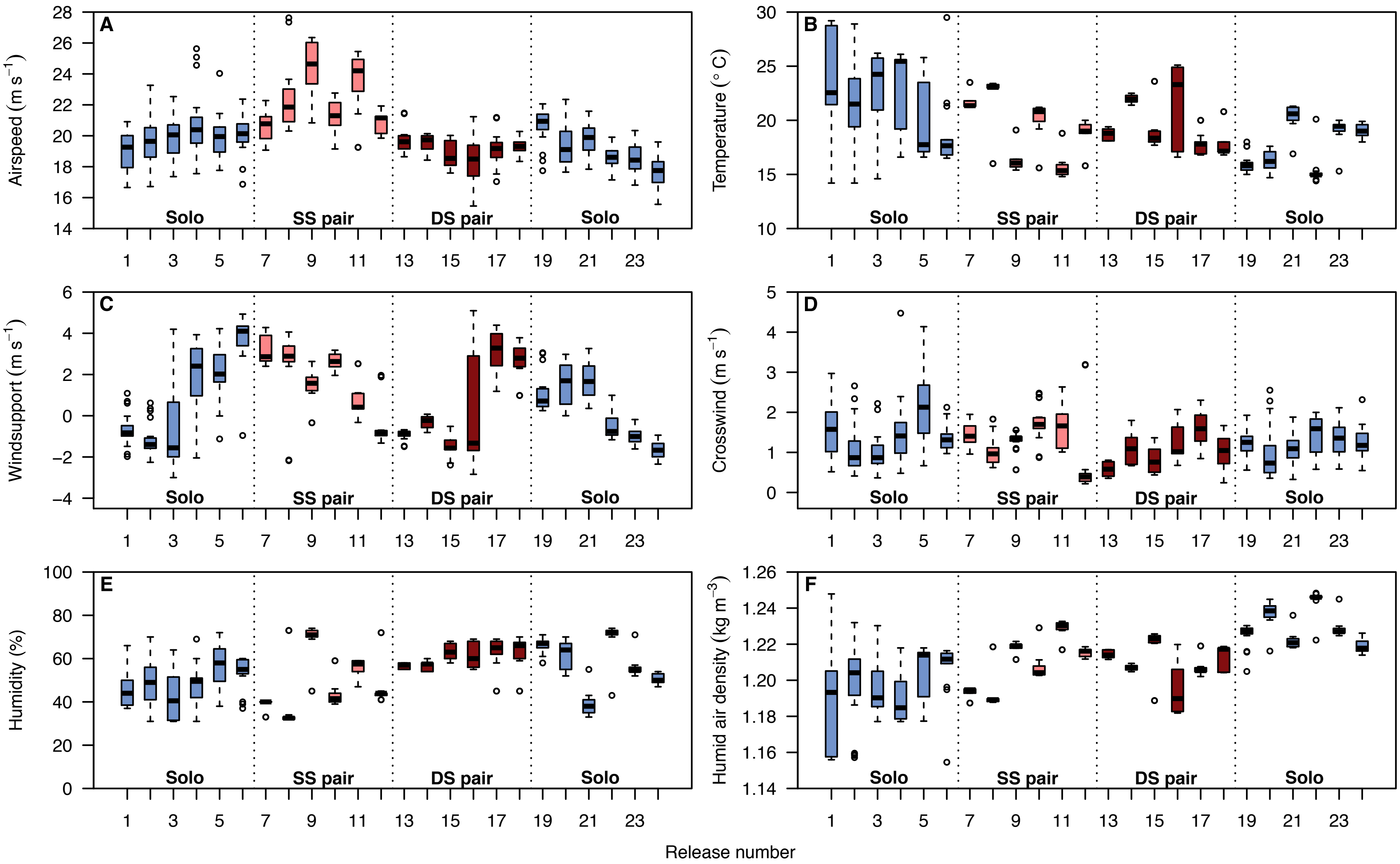


**Figure S2.** Raw data showing the results per release for the Bayesian hierarchical model covariates: (**A**) median airspeed (m s$^{-1}$), (**B**) median temperature (°C), (**C**) median wind support (m s$^{-1}$), (**D**) median crosswind, (**E**) median humidity (%), and (**F**) median humid air density (kg m$^{-3}$).

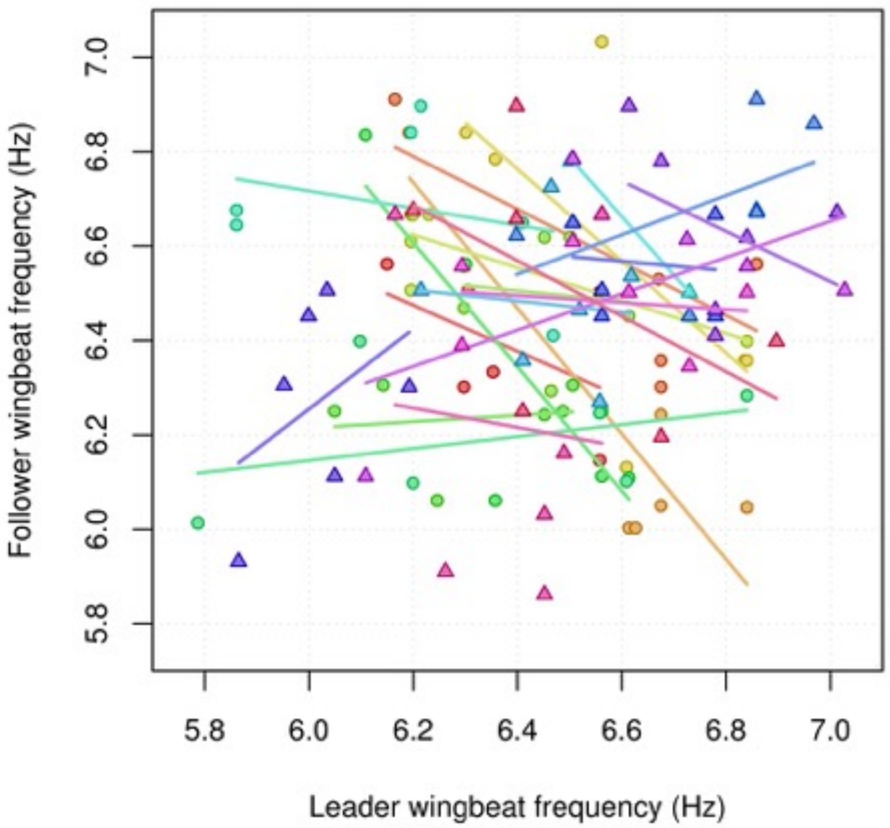


**Figure S3.** Median wingbeat frequency of following versus leading bird during each flight. The different coloured lines correspond to least squares regression lines fitted to the data from each pair, and the lack of any direct relationship indicates that the bird's wingbeats cannot be phase-locked through the flight.

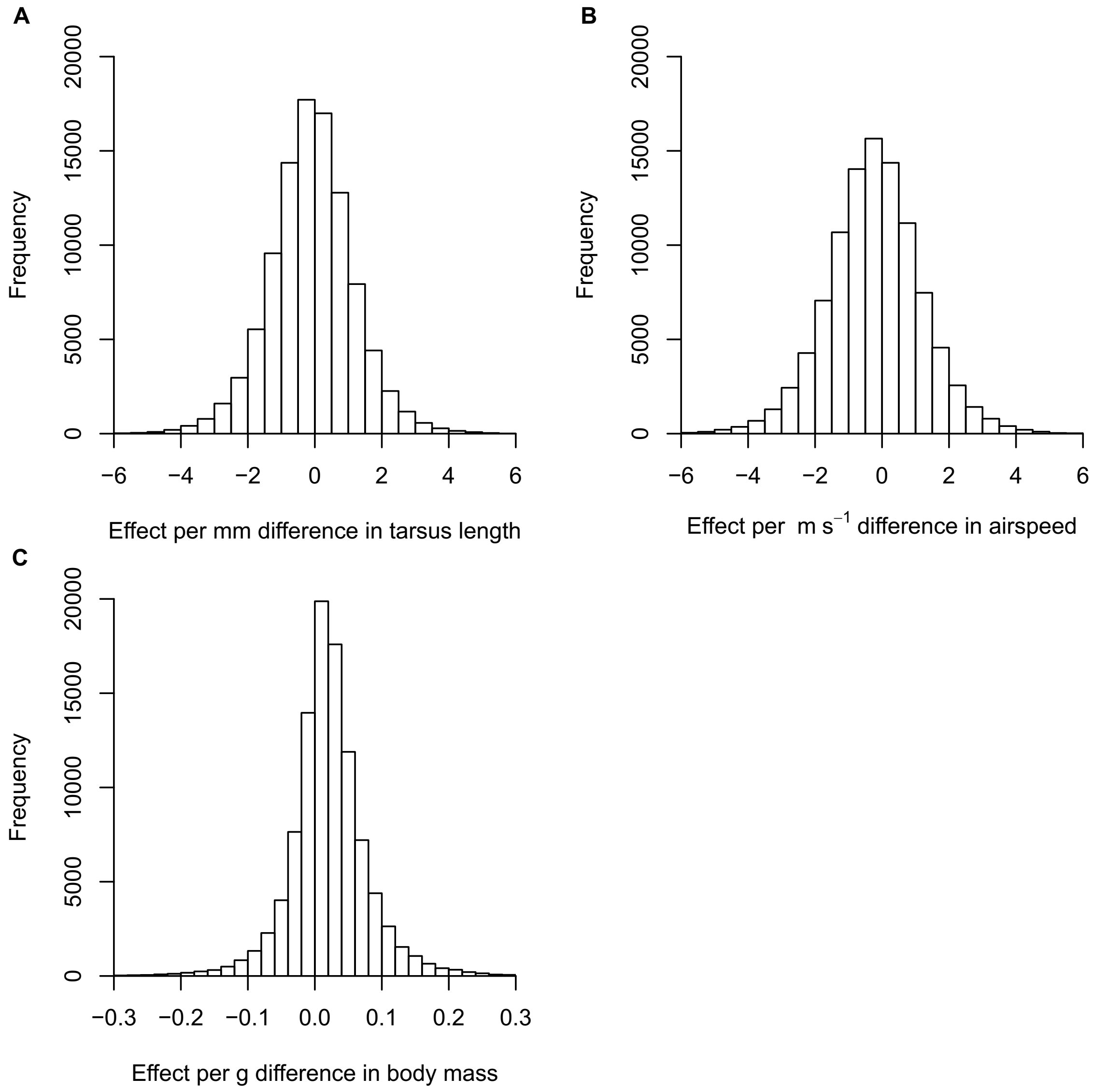


**Figure S4.** Probability of a bird leading per unit difference in (**A**) tarsus length, (**B**) median solo airspeed and (**C**) body mass.

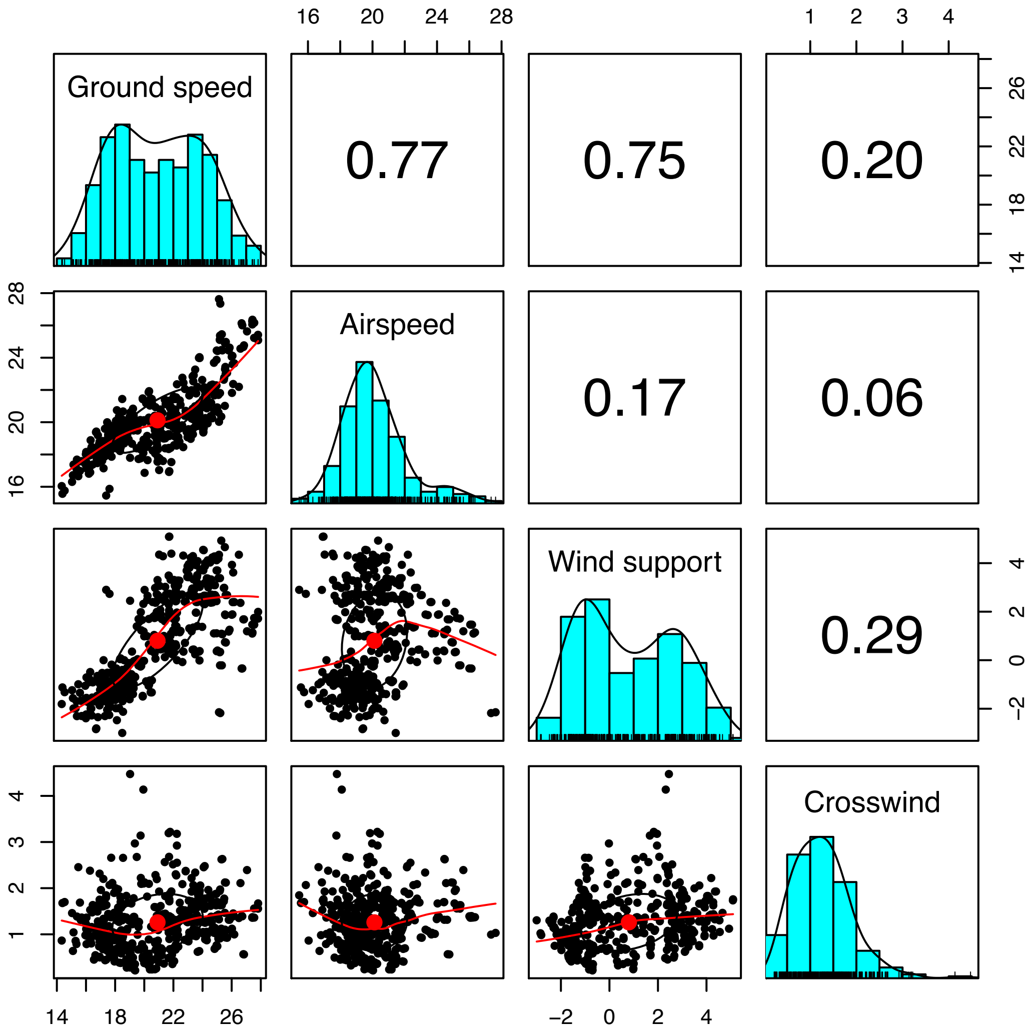


**Figure S5.** Relationship between ground speed, airspeed, wind support and crosswind.

**Table S1.** Results of the Bayesian hierarchical models. Corrected solo mean values refer to the mean of the median value from each individual, which has been corrected for the effects of airspeed, wind support, crosswind, temperature, humidity and air density according to the results of the model. Note: the covariates in this table are the difference per unit difference in the covariate.

| Variable | Corrected solo mean | Difference from solo for birds of the same tarsus length | | Difference per mm difference in tarsus length | | Covariates: Difference per unit difference in | | | | | | |
|---|---|---|---|---|---|---|---|---|---|---|---|---|
| | | Mean Δ [95% CrI] | % | Mean Δ [95% CrI] | % | Wind support (m $s^{-1}$) [95% CrI] max = 5.10 m $s^{-1}$ min = -3.00 m $s^{-1}$ | Crosswind (m $s^{-1}$) [95% CrI] max = 4.47 m $s^{-1}$ min = 0.31 m $s^{-1}$ | Temperature (ºC) [95% CrI] max = 29.4 ºC min = 14.2 ºC | Humidity (%) [95% CrI] max = 74 % min = 31 % | Air density (kg $m^{-3}$) [95% CrI] max = 1.25 kg $m^{-3}$ min = 1.15 kg $m^{-3}$ | Airspeed (m $s^{-1}$) [95% CrI] max = 27.63 m $s^{-1}$ min = 15.46 m $s^{-1}$ | Date of release |
| Median wingbeat frequency (Hz) | 5.48 | 1.00 [0.61, 1.38] | 18.24 | 0.01 [-0.13, 0.16] | 0.27 | 0.03 [-0.02, 0.07] | 0.02 [-0.01, 0.05] | 0.01 [-0.02, 0.04] | 0.00 [-0.01, 0.01] | -0.40 [-1.16, 0.36] | 0.01 [-0.01, 0.03] | Min: -0.40 Max: 0.43 |
| Standard deviation of wingbeat frequency (Hz) | 0.60 | -0.01 [-0.11, 0.09] | -2.16 | 0.02 [-0.15, 0.18] | 3.54 | -0.01 [-0.04, 0.01] | 0.00 [-0.01, 0.02] | 0.02 [0.00, 0.03] | 0.00 [0.00, 0.01] | -0.55 [-0.99, -0.11] | 0.02 [0.00, 0.03] | Min: -0.19 Max: 0.20 |
| Median downstroke phase duration (ms) | 47.67 | -3.53 [-8.35, 1.38] | -7.41 | -0.1 [-1.82, 1.56] | -0.21 | -0.07 [-0.69, 0.55] | 0.13 [-0.19, 0.44] | 0.01 [-0.17, 0.19] | 0 [-0.05, 0.04] | 0.87 [-2.83, 4.56] | -0.07 [-0.29, 0.15] | Min: -2.56 Max: 1.62 |
| Median upstroke phase duration (ms) | 135.76 | -27.99 [-35.61, -19.92] | -20.62 | 0.33 [-3.73, 4.58] | 0.24 | -0.16 [-1.39, 1.09] | 0.61 [0.04, 1.19] | -0.29 [-0.62, 0.03] | 0.01 [-0.08, 0.09] | -0.63 [-6.58, 5.32] | 0.4 [-0.02, 0.82] | Min: -7.47 Max: 7.53 |
| Median peak-to-peak DB acceleration (***g***) | 3.65 | 0.16 [-0.16, 0.47] | 4.35 | 0 [-0.33, 0.34] | -0.07 | 0.01 [-0.07, 0.09] | -0.02 [-0.07, 0.03] | -0.01 [-0.06, 0.03] | 0.00 [-0.01, 0.01] | -0.32 [-1.35, 0.71] | 0.03 [0.00, 0.07] | Min: -0.36 Max: 0.47 |
| Median DB amplitude (mm) | 20.68 | -4.65 [-6.21, -3.06] | -22.47 | 0.02 [-0.60, 0.57] | 0.08 | -0.09 [-0.34, 0.16] | 0.09 [-0.06, 0.24] | -0.09 [-0.20, 0.01] | 0 [-0.02, 0.02] | -0.33 [-2.77, 2.10] | 0.14 [0.04, 0.25] | Min: -1.61 Max: 2.29 |
| Airspeed (m $s^{-1}$) | 19.54 | 0.64 [0.08, 1.20] | 3.26 | -0.32 [-0.64, 0.05] | -1.62 | -0.11 [-0.31, 0.09] | -0.76 [-0.87, -0.65] | -0.03 [-0.11, 0.05] | 0.01 [0.00, 0.03] | 0.75 [-1.24, 2.75] | N/A | Min: -4.26 Max: 4.92 |
| Route accuracy | 0.80 | 0.06 [0.01, 0.10] | 6.93 | 0 [-0.02, 0.03] | 0.07 | -0.01 [-0.02, 0.00] | 0.02 [0.01, 0.03] | 0 [-0.01, 0.01] | 0.00 [0.00, 0.00] | -0.23 [-0.50, 0.04] | 0.01 [0.01, 0.02] | Min: -0.17 Max: 0.10 |

**Table S2.** A calculation of the total number of wingbeats that would have been used during solo and paired flight had the birds flown with the stated wingbeat frequency, mean route accuracy and at the stated mean airspeed in the absence of any wind support. We calculated the difference using the results of the Bayesian hierarchical models combined with the means of the solo flights after accounting for the effects of covariates as not all birds completed 100% of the paired flights together with one leader.

| | | Solo | Pair |
|---|---|---|---|
| Variables | Straight-line distance (m; excluding 200m take-off and landing) | 6660.58 | 6660.58 |
| | Mean corrected route accuracy | 0.80 | 0.86 |
| | Mean corrected airspeed (m $s^{-1}$) | 19.54 | 20.17 |
| | Mean corrected wingbeat frequency (Hz) | 5.48 | 6.48 |
| Calculations | Total flight distance (m) | 8295 | 7759 |
| | Total flight time (s) | 425 | 385 |
| | Total number of wingbeats | 2325 | 2490 |
| | Number of wingbeats difference | | +165.47 |
| | Percentage wingbeats difference (%) | | +7.12 |

**Table S3.** Bird and pairing information, including tarsus length (mm), body mass (g) and median solo airspeed (m $s^{-1}$).

| Individual information | | | | Similar-sized pairs | | | | Different-sized pairs | | | |
|---|---|---|---|---|---|---|---|---|---|---|---|
| Bird ID | Tarsus length (mm) | Body mass (g) | Median solo airspeed (m $s^{-1}$) | Partner ID | Δ Tarsus length (mm) | Δ Body mass (g) | Δ Median solo airspeed (m $s^{-1}$) | Partner ID | Δ Tarsus length (mm) | Δ Body mass (g) | Δ Median solo airspeed (m $s^{-1}$) |
| B01 | 32.4 | 452 | 19.0 | B82 | -0.9 | 21.0 | -0.1 | B07 | -2.8 | -39.0 | -1.0 |
| B07 | 35.2 | 491 | 20.0 | B87 | 0.3 | -28.0 | 0.6 | B01 | 2.8 | 39.0 | 1.0 |
| B61 | 35.2 | 478 | 19.5 | S25 | -0.8 | 15.0 | 0.5 | S87 | 3.8 | 48.0 | 1.9 |
| B65 | 35.5 | 454 | 18.3 | S84 | 1.0 | -31.0 | -0.5 | S13 | -0.7 | -75.0 | -1.4 |
| B67 | 36.0 | 474 | 20.7 | B80 | 0.0 | 3.0 | 0.6 | S05 | 1.3 | 7.0 | 0.4 |
| B75 | 35.4 | 461 | 18.9 | S06 | -0.2 | 17.0 | -1.5 | S30 | -0.7 | -33.0 | -1.5 |
| B80 | 36.0 | 471 | 20.1 | B67 | 0.0 | -3.0 | -0.6 | S12 | 1.5 | 2.0 | 1.0 |
| B82 | 33.3 | 431 | 19.1 | B01 | 0.9 | -21.0 | 0.1 | B87 | -1.6 | -88.0 | -0.3 |
| B84 | 33.8 | 425 | 20.0 | S87 | 2.4 | -5.0 | 2.5 | S25 | -2.2 | -38.0 | 1.0 |
| B87 | 34.9 | 519 | 19.5 | B07 | -0.3 | 28.0 | -0.6 | B82 | 1.6 | 88.0 | 0.3 |
| S03 | 36.1 | 493 | 19.3 | S30 | 0.0 | -1.0 | -1.1 | S06 | 0.5 | 49.0 | -1.1 |
| S05 | 34.7 | 467 | 20.3 | S12 | 0.2 | -2.0 | 1.1 | B67 | -1.3 | -7.0 | -0.4 |
| S06 | 35.6 | 444 | 20.4 | B75 | 0.2 | -17.0 | 1.5 | S03 | -0.5 | -49.0 | 1.1 |
| S12 | 34.5 | 469 | 19.1 | S05 | -0.2 | 2.0 | -1.1 | B80 | -1.5 | -2.0 | -1.0 |
| S13 | 36.2 | 529 | 19.7 | S27 | -0.1 | 6.0 | -0.5 | B65 | 0.7 | 75.0 | 1.4 |
| S25 | 36.0 | 463 | 19.0 | B61 | 0.8 | -15.0 | -0.5 | B84 | 2.2 | 38.0 | -1.0 |
| S27 | 36.3 | 523 | 20.2 | S13 | 0.1 | -6.0 | 0.5 | S84 | 1.8 | 38.0 | 1.4 |
| S30 | 36.1 | 494 | 20.4 | S03 | 0.0 | 1.0 | 1.1 | B75 | 0.7 | 33.0 | 1.5 |
| S84 | 34.5 | 485 | 18.8 | B65 | -1.0 | 31.0 | 0.5 | S27 | -1.8 | -38.0 | -1.4 |
| S87 | 31.4 | 430 | 17.6 | B84 | -2.4 | 5.0 | -2.5 | B61 | -3.8 | -48.0 | -1.9 |

**Table S4.** Model priors for the hyper-parameters of the Bayesian hierarchical model. The width of the prior distributions was chosen to be similar to that of the observed quantities. The hyper-parameters represent the population (i.e. across all birds) averages. The bird-level parameters are obtained by sampling from a normal distribution conditional on the hyper-parameters. See the Stan model for further information

| Variable | $\delta, \eta$ | $\chi$ | $\mu$ | $\gamma$ | $\sigma$ |
|---|---|---|---|---|---|
| Median wingbeat frequency (Hz) | normal(0,0.3) | normal(0,2) | normal(5.5,0.3) | normal(0,0.6) | cauchy(0,0.6) |
| Standard deviation of wingbeat frequency (Hz) | normal(0,0.1) | normal(0,2) | normal(0.6,0.1) | normal(0,0.4) | cauchy(0,0.4) |
| Median downstroke phase duration (ms) | normal(0,4.6) | normal(0,2) | normal(47.5,4.6) | normal(0,2.1) | cauchy(0,2.1) |
| Median upstroke phase duration (ms) | normal(0,11.1) | normal(0,2) | normal(135.9,11.1) | normal(0,3.3) | cauchy(0,3.3) |
| Median peak-to-peak DB acceleration (*g*) | normal(0,0.4) | normal(0,2) | normal(3.7,0.4) | normal(0,0.7) | cauchy(0,0.7) |
| Median DB amplitude (mm) | normal(0,2.2) | normal(0,2) | normal(20.7,2.2) | normal(0,1.5) | cauchy(0,1.5) |
| Airspeed (m $s^{-1}$) | normal(0,1.6) | normal(0,2) | normal(19.5,1.6) | normal(0,1.3) | cauchy(0,1.3) |
| Route accuracy | normal(0,0.1) | normal(0,2) | normal(0.8,0.1) | normal(0,0.3) | cauchy(0,0.3) |

**Supplementary material**

**Analysis of accelerometer output**

The purpose of this section is to provide a rigorous analysis of the oscillatory accelerations experienced by an accelerometer worn on a bird's back during flapping flight. Here, the axis of the accelerometer is assumed to be aligned dorsoventrally on the bird, and possible angular changes in the inclination of this axis with respect to the Earth are assumed to be sufficiently small or slow that the dorsoventral component of gravitational acceleration that the accelerometer experiences can be considered constant and hence neglected in an analysis of the oscillatory accelerations.

Under Newton's Second Law, an accelerometer placed at the centre of mass of an object experiences an acceleration $a(t)$ equal to the net external force on the object divided by its mass. If the accelerometer is placed away from the centre of mass, then the angular motion of the bird will also contribute to the sensed acceleration, but this effect may be neglected in non-manoeuvring flight. Under sinusoidal aerodynamic forcing, the accelerometer therefore experiences an aerodynamic acceleration:

$$\tilde{a}_A(t) = \frac{F \sin(2\pi f t)}{m} \quad \text{(Eq. S1)}$$

where $F$ is the aerodynamic forcing amplitude, $f$ is the wingbeat frequency, $t$ is time, and $m$ is the mass. Integrating twice with respect to time, this aerodynamic acceleration is associated with an oscillatory displacement of the centre of mass:

$$\tilde{d}_A(t) = -\frac{F \sin(2\pi f t)}{4\pi^2 f^2 m}$$

(Eq. S2)

Other things being equal, the aerodynamic displacement of the centre of mass therefore scales as $f^{-2}$, such that the 18% increase in wingbeat frequency that we measured between solo and paired flights (Table S1) could be expected to cause a 28% reduction in the amplitude of the aerodynamic body displacement. This conclusion holds provided that the increase in wingbeat frequency is not accompanied by any increase in the amplitude $F$ of the dorsoventral aerodynamic forcing, which is a reasonable assumption in sustained level flight, for which the time-averaged vertical force must always balance the weight of the bird (see main text). This being so, it follows that a bird can reduce the bobbing motion that its head or body experiences in flight simply by increasing its wingbeat frequency.

A bird is not a rigid body, however, and flapping its wings will cause its overall mass distribution to vary periodically. An accelerometer placed at any anatomically fixed point on the bird's body will therefore experience an additional inertial acceleration due to the displacement of the bird's body in relation to its overall centre of mass. The wings of a pigeon together comprise 13% of its overall mass, and their respective centres of mass move on radii of 0.07m from the shoulder joint [27]. Lowering the wings from the extreme dorsal to the extreme ventral position will therefore raise the bird's body by 0.018m relative to its overall centre of mass, calculated as 13% of the peak-to-peak displacement of the wing's centre of mass. Based on the wingtip kinematics presented by Tobalske & Dial [21], an inertial displacement nearer 80% of this extreme seems realistic at the 20ms$^{-1}$ airspeed that we observed (Table S1), bringing the expected peak-to-peak amplitude of the oscillatory inertial displacement $\tilde{d}_I(t)$ to approximately 0.015m. Since the amplitude of this displacement is independent of the wingbeat frequency, the presence of this inertial forcing is expected to attenuate the overall reduction of body displacement that would otherwise result from increasing the frequency of the aerodynamic forcing, which qualitatively speaking is what we observed (see below).

Neither the aerodynamic nor the inertial component of the total displacement is expected to be perfectly sinusoidal, but it is nevertheless informative to consider how their waveforms would be expected to combine if as a good first approximation they were. Approximating the dorsoventral motion of the wing's centre of mass as a simple sinusoid yields the following expression for the inertial displacement:

$$\tilde{d}_I(t) \approx 0.0075 \sin(2\pi f t + \varphi)$$

(Eq. S3)

where $\varphi$ is the phase of the motion with respect to the aerodynamic displacement $\tilde{d}_A(t)$ in Eq. S2, and where the numerical constant is half the peak-to-peak amplitude. Because the upward aerodynamic acceleration $\tilde{a}_A(t)$ of the accelerometer is expected to peak at mid-downstroke, its upward aerodynamic displacement $\tilde{d}_A(t)$ is expected to peak at mid-upstroke (see Eq. S2). Hence, since the upward inertial displacement of the accelerometer $\tilde{d}_I(t)$ peaks at the bottom of the downstroke, it follows that the inertial displacement $\tilde{d}_I(t)$ leads the aerodynamic displacement $\tilde{d}_A(t)$ by approximately 90° such that $\varphi = -\pi/2$.

The amplitude of a waveform combining two sinusoids with a 90° phase lag is just the square root of the sum of the squared amplitudes of the two sinusoids. Subtracting the squared peak-to-peak amplitude of the predicted displacement due to inertial forcing (0.015 m) from the squared peak-to-peak amplitude of the measured displacement (0.027 m in solo flight *versus* 0.022 m in paired flight; Table S1) and taking the square root, we therefore arrive at a robust empirical estimate of the peak-to-peak amplitude of the displacement due to aerodynamic forcing, which is 0.022m in solo flight and 0.016m in paired flight. The resulting empirical estimate of a 27% reduction in the peak-to-peak amplitude of the aerodynamic displacement between solo and paired flight coincides almost exactly with the 28% reduction that we predicted theoretically from the accompanying 18% increase in wingbeat frequency using Eq. S2 above. These two estimates are mathematically independent, and therefore provide a useful internal validation of our analysis. Moreover, these figures are modified only slightly if we assume that flying in pairs is associated with a 10% decrease in the wingbeat amplitude, as is argued in the main text. Multiplying our empirical estimates of the peak-to-peak amplitude of the aerodynamic displacement by the factor $4\pi^2 f^2$ in the denominator of Eq. S2, we arrive at an estimate of 2.7$g$ for the peak-to-peak amplitude of the acceleration due to aerodynamic forcing in solo flight, and 2.8$g$ in paired flight. This is consistent with our earlier argument that the amplitude $F$ of the dorsoventral aerodynamic forcing should have been similar in paired and solo flight, on the basis that the time-averaged vertical aerodynamic force must have balanced the weight of the bird under both flight conditions (see also the main text).

### Stan code for the Bayesian hierarchical model

Stan code for the Bayesian hierarchical model of wingbeat frequency. In the model, the bird-level parameters are sampled from a normal distribution conditional on the hyper-parameters, which represent the population averages. For example, $delta_i$ ~ N($delta^{\{top\}}$, $sigma\_delta^{\{top\}}$), where $delta_i$ is the uplift for bird $i$ of being in a pair, and $delta^{\{top\}}$ and $sigma\_delta^{\{top\}}$ represent the population mean uplift and the standard deviation of the individual birds' effects about this mean. Note that in the Stan model, we use a non-centred parameterisation to implement this model[45], which helps with sampling.

```
functions{
  // calculates mixing weight of solo frequencies of birds
  // based on difference in tarsus length
  real weighting(int bird_a, int bird_b, real[] chi_0, real[] chi, real[] tarsus){
    real omega;
    omega = inv_logit(chi_0[bird_a] + chi[bird_a] * fabs(tarsus[bird_a] - tarsus[bird_b]));
    return(omega);
  }
}

data{
  int N; // length of data
  real W[N]; // wingbeat frequency
  int nBirds; // number of birds

  // indices
  int bird1[N]; // identity of first bird in pair
  int bird2[N]; // identity of second bird in pair -- can be same as first

  // covariates
  real wind1[N];
  real wind2[N];
  real temp[N];
  real density[N];
  real humidity[N];
  real airspeed[N];

  // matrix covariates
  int<lower=1> K;
  matrix[N, K] date;

  // size covariates
  real tarsus[nBirds];
}

parameters{
  // bird-specific
  real mu[nBirds]; // mean wingbeat for each bird
  real delta_0_raw[nBirds]; // uplift from being in a pair
  real eta_raw[nBirds]; // effect of size difference on wingbeat

  // top-level
```

```
  real delta_0_top;
  real eta_top;
  real<lower=0> sigma_delta_0_top;
  real<lower=0> sigma_eta_top;

  // covariate specific
  real gamma1;
  real gamma2;
  real gamma3;
  real gamma4;
  real gamma5;
  real gamma6;

  // matrix covariate
  vector[K] beta;

  // other
  real<lower=0> sigma;

  // Leading parameter
  real chi_raw[nBirds];
  real chi_0_raw[nBirds];
  real chi_top;
  real chi_0_top;
  real<lower=0> chi_sigma_top;
  real<lower=0> chi_0_sigma_top;
}

transformed parameters{
  real chi[nBirds];
  real chi_0[nBirds];
  real delta_0[nBirds]; // uplift from being in a pair
  real eta[nBirds]; // effect of size difference on wingbeat

  for(i in 1:nBirds){
    chi_0[i] = chi_0_top + chi_0_raw[i] * chi_0_sigma_top;
    chi[i] = chi_top + chi_raw[i] * chi_sigma_top;
    delta_0[i] = delta_0_top + delta_0_raw[i] * sigma_delta_0_top;
    eta[i] = eta_top + eta_raw[i] * sigma_eta_top;
    }
}

model{
  real omega;
  for(i in 1:N){
    real aMean;
    if(bird1[i] == bird2[i]){ // solo
      aMean = 0.0;
    else{
      aMean = delta_0[bird1[i]] + eta[bird1[i]] * fabs(tarsus[bird1[i]] - tarsus[bird2[i]]);
```

```
      }
      // mixing probability
      omega = weighting(bird1[i], bird2[i], chi_0, chi, tarsus);

      // likelihood
      W[i] ~ normal(aMean + omega * mu[bird1[i]] + (1 - omega) * mu[bird2[i]]
                  + wind1[i] * gamma1 + wind2[i] * gamma2 + temp[i] * gamma3
                  + density[i] * gamma4 + humidity[i] * gamma5 + airspeed[i] * gamma6
                  + date[i] * beta, sigma);
    }

    // priors
    chi_0_raw ~ normal(0, 1);
    chi_raw ~ normal(0, 1);
    delta_0_raw ~ normal(0, 1);
    eta_raw ~ normal(0, 1);
    mu ~ normal(5.5, 0.3); // change mean to be similar to that observed + sd change
    gamma1 ~ normal(0, 0.6);
    gamma2 ~ normal(0, 0.6);
    gamma3 ~ normal(0, 0.6);
    gamma4 ~ normal(0, 0.6);
    gamma5 ~ normal(0, 0.6);
    gamma6 ~ normal(0, 0.6);
    beta ~ normal(0, 0.6);
    sigma ~ cauchy(0,0.6);

    // hyper-priors
    chi_0_top ~ normal(0, 2);
    chi_top ~ normal(0, 2);
    chi_sigma_top ~ cauchy(0, 2);
    chi_0_sigma_top ~ cauchy(0, 2);
    delta_0_top ~ normal(0, 0.3);
    eta_top ~ normal(0, 0.3);
    sigma_delta_0_top ~ cauchy(0, 0.6);
    sigma_eta_top ~ cauchy(0, 0.6);
  }

  generated quantities{
    real delta_0_av;
    real eta_av;
    real chi_av;
    real chi_0_av;

    delta_0_av = normal_rng(delta_0_top, sigma_delta_0_top);
    eta_av = normal_rng(eta_top, sigma_eta_top);
    chi_av = normal_rng(chi_top, chi_sigma_top);
    chi_0_av = normal_rng(chi_0_top, chi_0_sigma_top);
  }

```

Inference for Stan model: wingbeat_continuous_pred_MEDIAN_FPS_wDate.
8 chains, each with iter=25000; warmup=12500; thin=1;
post-warmup draws per chain=12500, total post-warmup draws=1e+05.

| | mean | se_mean | sd | 2.5% | 25% | 50% | 75% | 97.5% | n_eff | Rhat |
|---|---|---|---|---|---|---|---|---|---|---|
| mu[1] | 5.48 | 0.00 | 0.08 | 5.32 | 5.42 | 5.48 | 5.54 | 5.64 | 12695 | 1 |
| mu[2] | 5.33 | 0.00 | 0.08 | 5.16 | 5.27 | 5.33 | 5.38 | 5.49 | 13119 | 1 |
| mu[3] | 5.50 | 0.00 | 0.08 | 5.33 | 5.44 | 5.50 | 5.55 | 5.67 | 12531 | 1 |
| mu[4] | 5.69 | 0.00 | 0.09 | 5.52 | 5.63 | 5.69 | 5.74 | 5.86 | 13599 | 1 |
| mu[5] | 5.09 | 0.00 | 0.09 | 4.92 | 5.03 | 5.09 | 5.14 | 5.25 | 13127 | 1 |
| mu[6] | 5.43 | 0.00 | 0.08 | 5.26 | 5.37 | 5.43 | 5.49 | 5.60 | 13183 | 1 |
| mu[7] | 5.77 | 0.00 | 0.08 | 5.60 | 5.71 | 5.77 | 5.82 | 5.93 | 13079 | 1 |
| mu[8] | 5.18 | 0.00 | 0.08 | 5.01 | 5.12 | 5.18 | 5.23 | 5.34 | 12837 | 1 |
| mu[9] | 5.47 | 0.00 | 0.09 | 5.30 | 5.41 | 5.47 | 5.53 | 5.64 | 13580 | 1 |
| mu[10] | 5.65 | 0.00 | 0.08 | 5.49 | 5.60 | 5.65 | 5.71 | 5.82 | 13129 | 1 |
| mu[11] | 5.40 | 0.00 | 0.08 | 5.23 | 5.34 | 5.40 | 5.46 | 5.57 | 12668 | 1 |
| mu[12] | 5.37 | 0.00 | 0.09 | 5.20 | 5.31 | 5.37 | 5.43 | 5.55 | 13877 | 1 |
| mu[13] | 5.63 | 0.00 | 0.09 | 5.46 | 5.58 | 5.63 | 5.69 | 5.81 | 13688 | 1 |
| mu[14] | 5.45 | 0.00 | 0.08 | 5.28 | 5.39 | 5.45 | 5.51 | 5.61 | 12893 | 1 |
| mu[15] | 5.35 | 0.00 | 0.09 | 5.17 | 5.29 | 5.35 | 5.42 | 5.53 | 15439 | 1 |
| mu[16] | 5.65 | 0.00 | 0.09 | 5.48 | 5.59 | 5.65 | 5.71 | 5.82 | 13871 | 1 |
| mu[17] | 5.52 | 0.00 | 0.10 | 5.33 | 5.45 | 5.52 | 5.58 | 5.71 | 16426 | 1 |
| mu[18] | 5.76 | 0.00 | 0.08 | 5.60 | 5.70 | 5.76 | 5.82 | 5.92 | 12997 | 1 |
| mu[19] | 5.73 | 0.00 | 0.08 | 5.57 | 5.68 | 5.73 | 5.79 | 5.90 | 12928 | 1 |
| mu[20] | 5.48 | 0.00 | 0.09 | 5.30 | 5.42 | 5.48 | 5.54 | 5.65 | 14507 | 1 |
| delta_0_raw[1] | 0.63 | 0.00 | 0.72 | -0.80 | 0.15 | 0.63 | 1.10 | 2.02 | 39555 | 1 |
| delta_0_raw[2] | 0.29 | 0.00 | 0.57 | -0.88 | -0.08 | 0.29 | 0.66 | 1.40 | 60260 | 1 |
| delta_0_raw[3] | -0.98 | 0.00 | 0.58 | -2.16 | -1.35 | -0.96 | -0.59 | 0.14 | 61267 | 1 |
| delta_0_raw[4] | -0.48 | 0.00 | 0.55 | -1.61 | -0.82 | -0.46 | -0.12 | 0.57 | 54396 | 1 |
| delta_0_raw[5] | -0.20 | 0.00 | 0.61 | -1.49 | -0.58 | -0.17 | 0.21 | 0.94 | 50238 | 1 |
| delta_0_raw[6] | -0.10 | 0.00 | 0.55 | -1.21 | -0.46 | -0.09 | 0.26 | 0.97 | 55738 | 1 |
| delta_0_raw[7] | 0.37 | 0.00 | 0.58 | -0.79 | -0.01 | 0.37 | 0.75 | 1.49 | 53970 | 1 |
| delta_0_raw[8] | -0.48 | 0.00 | 0.69 | -1.81 | -0.94 | -0.49 | -0.02 | 0.90 | 61057 | 1 |
| delta_0_raw[9] | -1.18 | 0.00 | 0.73 | -2.54 | -1.66 | -1.21 | -0.74 | 0.40 | 48195 | 1 |
| delta_0_raw[10] | 1.48 | 0.00 | 0.59 | 0.35 | 1.09 | 1.48 | 1.86 | 2.67 | 61922 | 1 |
| delta_0_raw[11] | 0.58 | 0.00 | 0.55 | -0.51 | 0.23 | 0.58 | 0.94 | 1.65 | 56837 | 1 |
| delta_0_raw[12] | -0.50 | 0.00 | 0.53 | -1.61 | -0.84 | -0.48 | -0.14 | 0.48 | 55758 | 1 |
| delta_0_raw[13] | -0.65 | 0.00 | 0.57 | -1.80 | -1.01 | -0.63 | -0.27 | 0.44 | 46209 | 1 |
| delta_0_raw[14] | 1.23 | 0.00 | 0.53 | 0.24 | 0.87 | 1.21 | 1.57 | 2.31 | 66061 | 1 |
| delta_0_raw[15] | 0.81 | 0.00 | 0.57 | -0.31 | 0.44 | 0.81 | 1.18 | 1.94 | 57962 | 1 |
| delta_0_raw[16] | 0.50 | 0.00 | 0.58 | -0.61 | 0.13 | 0.49 | 0.86 | 1.70 | 59497 | 1 |
| delta_0_raw[17] | -0.17 | 0.00 | 0.54 | -1.27 | -0.52 | -0.16 | 0.19 | 0.87 | 62711 | 1 |
| delta_0_raw[18] | 0.72 | 0.00 | 0.53 | -0.29 | 0.37 | 0.71 | 1.06 | 1.79 | 61395 | 1 |
| delta_0_raw[19] | -1.10 | 0.00 | 0.59 | -2.30 | -1.48 | -1.10 | -0.72 | 0.05 | 61094 | 1 |
| delta_0_raw[20] | 1.11 | 0.00 | 0.71 | -0.35 | 0.66 | 1.12 | 1.57 | 2.49 | 64988 | 1 |
| eta_raw[1] | 0.68 | 0.00 | 0.85 | -1.11 | 0.15 | 0.70 | 1.24 | 2.30 | 65991 | 1 |
| eta_raw[2] | 0.15 | 0.00 | 0.81 | -1.51 | -0.36 | 0.16 | 0.68 | 1.75 | 79650 | 1 |
| eta_raw[3] | -0.21 | 0.00 | 0.77 | -1.73 | -0.70 | -0.22 | 0.26 | 1.36 | 67529 | 1 |
| eta_raw[4] | -0.07 | 0.00 | 0.96 | -1.94 | -0.72 | -0.07 | 0.58 | 1.84 | 100000 | 1 |

| | | | | | | | | | | | |
|---|---|---|---|---|---|---|---|---|---|---|---|
| 382 | eta_raw[5] | -0.42 | 0.00 | 0.92 | -2.20 | -1.04 | -0.44 | 0.18 | 1.44 | 100000 | 1 |
| 383 | eta_raw[6] | 0.31 | 0.00 | 1.01 | -1.68 | -0.37 | 0.31 | 0.99 | 2.27 | 100000 | 1 |
| 384 | eta_raw[7] | -0.22 | 0.00 | 0.94 | -2.03 | -0.86 | -0.23 | 0.41 | 1.67 | 100000 | 1 |
| 385 | eta_raw[8] | 0.10 | 0.00 | 0.97 | -1.80 | -0.56 | 0.09 | 0.75 | 2.01 | 100000 | 1 |
| 386 | eta_raw[9] | -0.56 | 0.00 | 0.93 | -2.29 | -1.19 | -0.60 | 0.04 | 1.37 | 56750 | 1 |
| 387 | eta_raw[10] | -0.12 | 0.00 | 1.00 | -2.07 | -0.80 | -0.11 | 0.56 | 1.81 | 100000 | 1 |
| 388 | eta_raw[11] | 0.24 | 0.00 | 0.99 | -1.72 | -0.43 | 0.24 | 0.90 | 2.18 | 100000 | 1 |
| 389 | eta_raw[12] | 0.14 | 0.00 | 0.92 | -1.66 | -0.48 | 0.14 | 0.75 | 1.96 | 100000 | 1 |
| 390 | eta_raw[13] | 0.16 | 0.00 | 1.00 | -1.80 | -0.51 | 0.17 | 0.83 | 2.11 | 100000 | 1 |
| 391 | eta_raw[14] | 0.16 | 0.00 | 0.96 | -1.76 | -0.48 | 0.18 | 0.82 | 2.00 | 100000 | 1 |
| 392 | eta_raw[15] | 0.02 | 0.00 | 0.97 | -1.89 | -0.63 | 0.02 | 0.68 | 1.92 | 100000 | 1 |
| 393 | eta_raw[16] | -0.47 | 0.00 | 0.90 | -2.18 | -1.07 | -0.50 | 0.10 | 1.37 | 73506 | 1 |
| 394 | eta_raw[17] | 0.23 | 0.00 | 0.91 | -1.60 | -0.37 | 0.24 | 0.84 | 1.99 | 100000 | 1 |
| 395 | eta_raw[18] | -0.01 | 0.00 | 0.95 | -1.88 | -0.65 | -0.01 | 0.63 | 1.88 | 100000 | 1 |
| 396 | eta_raw[19] | -0.29 | 0.00 | 0.95 | -2.12 | -0.94 | -0.31 | 0.33 | 1.63 | 100000 | 1 |
| 397 | eta_raw[20] | 0.20 | 0.00 | 0.82 | -1.49 | -0.33 | 0.21 | 0.73 | 1.80 | 65524 | 1 |
| 398 | delta_0_top | 1.00 | 0.00 | 0.08 | 0.85 | 0.95 | 1.00 | 1.05 | 1.14 | 37710 | 1 |
| 399 | eta_top | 0.02 | 0.00 | 0.04 | -0.06 | -0.01 | 0.01 | 0.04 | 0.09 | 43410 | 1 |
| 400 | sigma_delta_0_top | 0.17 | 0.00 | 0.04 | 0.10 | 0.14 | 0.17 | 0.20 | 0.27 | 28315 | 1 |
| 401 | sigma_eta_top | 0.05 | 0.00 | 0.03 | 0.00 | 0.02 | 0.04 | 0.07 | 0.13 | 19474 | 1 |
| 402 | gamma1 | 0.02 | 0.00 | 0.02 | -0.01 | 0.01 | 0.02 | 0.03 | 0.05 | 32409 | 1 |
| 403 | gamma2 | 0.03 | 0.00 | 0.02 | -0.02 | 0.01 | 0.03 | 0.04 | 0.07 | 83371 | 1 |
| 404 | gamma3 | 0.01 | 0.00 | 0.02 | -0.02 | 0.00 | 0.01 | 0.02 | 0.04 | 21735 | 1 |
| 405 | gamma4 | -0.40 | 0.00 | 0.38 | -1.16 | -0.66 | -0.40 | -0.14 | 0.36 | 24497 | 1 |
| 406 | gamma5 | 0.00 | 0.00 | 0.00 | -0.01 | 0.00 | 0.00 | 0.00 | 0.01 | 23728 | 1 |
| 407 | gamma6 | 0.01 | 0.00 | 0.01 | -0.01 | 0.00 | 0.01 | 0.01 | 0.03 | 34661 | 1 |
| 408 | beta[1] | -0.18 | 0.00 | 0.22 | -0.61 | -0.33 | -0.18 | -0.03 | 0.26 | 23083 | 1 |
| 409 | beta[2] | -0.07 | 0.00 | 0.17 | -0.41 | -0.19 | -0.07 | 0.05 | 0.27 | 14841 | 1 |
| 410 | beta[3] | -0.04 | 0.00 | 0.17 | -0.36 | -0.15 | -0.04 | 0.07 | 0.28 | 12868 | 1 |
| 411 | beta[4] | -0.11 | 0.00 | 0.15 | -0.40 | -0.21 | -0.11 | -0.01 | 0.19 | 10082 | 1 |
| 412 | beta[5] | -0.37 | 0.00 | 0.14 | -0.65 | -0.46 | -0.37 | -0.27 | -0.09 | 9383 | 1 |
| 413 | beta[6] | -0.30 | 0.00 | 0.15 | -0.58 | -0.39 | -0.30 | -0.20 | -0.01 | 9556 | 1 |
| 414 | beta[7] | -0.40 | 0.00 | 0.16 | -0.73 | -0.51 | -0.40 | -0.29 | -0.08 | 11510 | 1 |
| 415 | beta[8] | -0.15 | 0.00 | 0.15 | -0.45 | -0.25 | -0.15 | -0.05 | 0.15 | 10360 | 1 |
| 416 | beta[9] | 0.06 | 0.00 | 0.15 | -0.23 | -0.04 | 0.06 | 0.16 | 0.35 | 9876 | 1 |
| 417 | beta[10] | 0.16 | 0.00 | 0.18 | -0.20 | 0.04 | 0.16 | 0.28 | 0.52 | 11046 | 1 |
| 418 | beta[11] | 0.10 | 0.00 | 0.14 | -0.17 | 0.01 | 0.10 | 0.20 | 0.37 | 8260 | 1 |
| 419 | beta[12] | -0.18 | 0.00 | 0.17 | -0.50 | -0.29 | -0.18 | -0.06 | 0.15 | 9976 | 1 |
| 420 | beta[13] | -0.14 | 0.00 | 0.13 | -0.41 | -0.23 | -0.14 | -0.05 | 0.12 | 8407 | 1 |
| 421 | beta[14] | -0.16 | 0.00 | 0.13 | -0.41 | -0.24 | -0.16 | -0.07 | 0.09 | 7335 | 1 |
| 422 | beta[15] | -0.08 | 0.00 | 0.14 | -0.34 | -0.17 | -0.08 | 0.02 | 0.20 | 8157 | 1 |
| 423 | beta[16] | -0.07 | 0.00 | 0.13 | -0.32 | -0.16 | -0.07 | 0.02 | 0.19 | 7647 | 1 |
| 424 | beta[17] | -0.02 | 0.00 | 0.15 | -0.31 | -0.12 | -0.02 | 0.08 | 0.27 | 9042 | 1 |
| 425 | beta[18] | 0.13 | 0.00 | 0.13 | -0.13 | 0.03 | 0.12 | 0.21 | 0.38 | 8155 | 1 |
| 426 | beta[19] | 0.01 | 0.00 | 0.19 | -0.35 | -0.11 | 0.01 | 0.14 | 0.38 | 15475 | 1 |
| 427 | beta[20] | 0.21 | 0.00 | 0.13 | -0.04 | 0.13 | 0.21 | 0.30 | 0.47 | 7874 | 1 |
| 428 | beta[21] | 0.13 | 0.00 | 0.14 | -0.14 | 0.03 | 0.13 | 0.22 | 0.40 | 8036 | 1 |
| 429 | beta[22] | 0.29 | 0.00 | 0.13 | 0.03 | 0.20 | 0.29 | 0.38 | 0.55 | 8156 | 1 |
| 430 | beta[23] | 0.05 | 0.00 | 0.17 | -0.29 | -0.07 | 0.05 | 0.16 | 0.38 | 10427 | 1 |
| 431 | beta[24] | -0.06 | 0.00 | 0.13 | -0.31 | -0.15 | -0.06 | 0.03 | 0.20 | 7775 | 1 |

| | | | | | | | | | | | |
|---|---|---|---|---|---|---|---|---|---|---|---|
| 432 | beta[25] | -0.08 | 0.00 | 0.13 | -0.33 | -0.16 | -0.08 | 0.01 | 0.18 | 7561 | 1 |
| 433 | beta[26] | 0.14 | 0.00 | 0.13 | -0.10 | 0.06 | 0.14 | 0.23 | 0.38 | 7379 | 1 |
| 434 | beta[27] | 0.11 | 0.00 | 0.12 | -0.13 | 0.03 | 0.11 | 0.19 | 0.35 | 6950 | 1 |
| 435 | beta[28] | 0.23 | 0.00 | 0.13 | -0.01 | 0.15 | 0.23 | 0.32 | 0.48 | 7322 | 1 |
| 436 | beta[29] | 0.43 | 0.00 | 0.14 | 0.16 | 0.34 | 0.43 | 0.52 | 0.70 | 8657 | 1 |
| 437 | sigma | 0.18 | 0.00 | 0.01 | 0.17 | 0.18 | 0.18 | 0.19 | 0.20 | 100000 | 1 |
| 438 | chi_raw[1] | 0.23 | 0.00 | 1.00 | -1.80 | -0.43 | 0.28 | 0.90 | 2.12 | 61955 | 1 |
| 439 | chi_raw[2] | -0.02 | 0.00 | 0.98 | -1.93 | -0.68 | -0.03 | 0.63 | 1.93 | 100000 | 1 |
| 440 | chi_raw[3] | 0.19 | 0.00 | 0.95 | -1.72 | -0.43 | 0.20 | 0.82 | 2.06 | 100000 | 1 |
| 441 | chi_raw[4] | -0.32 | 0.00 | 0.95 | -2.11 | -0.95 | -0.36 | 0.29 | 1.66 | 78330 | 1 |
| 442 | chi_raw[5] | 0.53 | 0.00 | 0.86 | -1.26 | 0.00 | 0.53 | 1.07 | 2.22 | 65970 | 1 |
| 443 | chi_raw[6] | -0.81 | 0.00 | 0.95 | -2.52 | -1.44 | -0.88 | -0.25 | 1.30 | 58717 | 1 |
| 444 | chi_raw[7] | -0.32 | 0.00 | 0.93 | -2.10 | -0.92 | -0.36 | 0.24 | 1.66 | 58195 | 1 |
| 445 | chi_raw[8] | -0.48 | 0.00 | 0.81 | -2.05 | -0.97 | -0.50 | -0.04 | 1.31 | 54191 | 1 |
| 446 | chi_raw[9] | 0.71 | 0.00 | 0.86 | -1.15 | 0.20 | 0.72 | 1.26 | 2.38 | 65672 | 1 |
| 447 | chi_raw[10] | -0.53 | 0.00 | 0.81 | -2.07 | -1.02 | -0.56 | -0.10 | 1.28 | 43879 | 1 |
| 448 | chi_raw[11] | -0.29 | 0.00 | 1.01 | -2.19 | -0.99 | -0.33 | 0.38 | 1.75 | 100000 | 1 |
| 449 | chi_raw[12] | 0.46 | 0.00 | 0.86 | -1.30 | -0.08 | 0.45 | 1.01 | 2.18 | 100000 | 1 |
| 450 | chi_raw[13] | 0.24 | 0.00 | 0.94 | -1.66 | -0.38 | 0.25 | 0.86 | 2.06 | 100000 | 1 |
| 451 | chi_raw[14] | 0.04 | 0.00 | 0.93 | -1.79 | -0.56 | 0.02 | 0.63 | 1.91 | 72609 | 1 |
| 452 | chi_raw[15] | 0.16 | 0.00 | 0.92 | -1.65 | -0.44 | 0.15 | 0.77 | 2.00 | 100000 | 1 |
| 453 | chi_raw[16] | -0.19 | 0.00 | 0.92 | -1.93 | -0.79 | -0.26 | 0.37 | 1.77 | 70294 | 1 |
| 454 | chi_raw[17] | -0.27 | 0.00 | 0.96 | -2.11 | -0.91 | -0.31 | 0.35 | 1.72 | 100000 | 1 |
| 455 | chi_raw[18] | 0.14 | 0.00 | 0.91 | -1.63 | -0.46 | 0.12 | 0.73 | 1.98 | 100000 | 1 |
| 456 | chi_raw[19] | 0.02 | 0.00 | 1.00 | -1.97 | -0.65 | 0.03 | 0.69 | 1.98 | 100000 | 1 |
| 457 | chi_raw[20] | 0.13 | 0.00 | 1.01 | -1.89 | -0.55 | 0.16 | 0.81 | 2.04 | 100000 | 1 |
| 458 | chi_0_raw[1] | -0.02 | 0.00 | 1.00 | -1.95 | -0.69 | -0.03 | 0.66 | 1.94 | 100000 | 1 |
| 459 | chi_0_raw[2] | 0.04 | 0.00 | 0.97 | -1.86 | -0.61 | 0.04 | 0.69 | 1.97 | 100000 | 1 |
| 460 | chi_0_raw[3] | 0.11 | 0.00 | 0.98 | -1.82 | -0.55 | 0.11 | 0.77 | 2.03 | 100000 | 1 |
| 461 | chi_0_raw[4] | -0.20 | 0.00 | 0.98 | -2.10 | -0.86 | -0.22 | 0.45 | 1.76 | 100000 | 1 |
| 462 | chi_0_raw[5] | 0.05 | 0.00 | 0.98 | -1.85 | -0.62 | 0.03 | 0.70 | 2.01 | 100000 | 1 |
| 463 | chi_0_raw[6] | -0.25 | 0.00 | 0.97 | -2.13 | -0.90 | -0.26 | 0.40 | 1.71 | 100000 | 1 |
| 464 | chi_0_raw[7] | 0.23 | 0.00 | 0.92 | -1.58 | -0.37 | 0.23 | 0.84 | 2.05 | 100000 | 1 |
| 465 | chi_0_raw[8] | -0.01 | 0.00 | 0.96 | -1.90 | -0.65 | -0.01 | 0.63 | 1.88 | 100000 | 1 |
| 466 | chi_0_raw[9] | 0.21 | 0.00 | 1.01 | -1.80 | -0.47 | 0.21 | 0.90 | 2.18 | 100000 | 1 |
| 467 | chi_0_raw[10] | 0.17 | 0.00 | 0.92 | -1.66 | -0.43 | 0.17 | 0.77 | 1.97 | 100000 | 1 |
| 468 | chi_0_raw[11] | 0.00 | 0.00 | 0.97 | -1.89 | -0.65 | -0.01 | 0.64 | 1.92 | 100000 | 1 |
| 469 | chi_0_raw[12] | 0.23 | 0.00 | 0.97 | -1.70 | -0.42 | 0.24 | 0.89 | 2.12 | 100000 | 1 |
| 470 | chi_0_raw[13] | -0.02 | 0.00 | 1.03 | -2.04 | -0.72 | -0.01 | 0.69 | 1.99 | 100000 | 1 |
| 471 | chi_0_raw[14] | 0.06 | 0.00 | 0.97 | -1.85 | -0.58 | 0.07 | 0.71 | 1.96 | 100000 | 1 |
| 472 | chi_0_raw[15] | 0.00 | 0.00 | 1.01 | -1.97 | -0.69 | 0.00 | 0.68 | 1.97 | 100000 | 1 |
| 473 | chi_0_raw[16] | 0.10 | 0.00 | 0.97 | -1.83 | -0.55 | 0.11 | 0.76 | 2.00 | 100000 | 1 |
| 474 | chi_0_raw[17] | -0.14 | 0.00 | 0.99 | -2.05 | -0.80 | -0.15 | 0.52 | 1.82 | 100000 | 1 |
| 475 | chi_0_raw[18] | 0.22 | 0.00 | 0.95 | -1.66 | -0.40 | 0.22 | 0.85 | 2.08 | 100000 | 1 |
| 476 | chi_0_raw[19] | -0.01 | 0.00 | 1.00 | -1.98 | -0.69 | -0.01 | 0.66 | 1.97 | 100000 | 1 |
| 477 | chi_0_raw[20] | 0.02 | 0.00 | 1.01 | -1.96 | -0.66 | 0.02 | 0.71 | 1.98 | 100000 | 1 |
| 478 | chi_top | -0.91 | 0.01 | 1.30 | -3.22 | -1.72 | -1.03 | -0.23 | 2.11 | 37663 | 1 |
| 479 | chi_0_top | 2.46 | 0.01 | 1.08 | 0.62 | 1.74 | 2.36 | 3.10 | 4.85 | 44115 | 1 |
| 480 | chi_sigma_top | 3.31 | 0.05 | 5.45 | 0.17 | 1.25 | 2.28 | 3.98 | 12.03 | 14355 | 1 |
| 481 | chi_0_sigma_top | 1.38 | 0.03 | 3.73 | 0.04 | 0.41 | 0.88 | 1.63 | 5.11 | 16724 | 1 |

| | | | | | | | | | | | |
|---|---|---|---|---|---|---|---|---|---|---|---|
| 482 | chi[1] | -0.08 | 0.05 | 7.55 | -7.95 | -2.02 | -0.49 | 1.28 | 10.20 | 20061 | 1 |
| 483 | chi[2] | -0.81 | 0.04 | 6.21 | -8.84 | -2.64 | -1.19 | 0.43 | 9.19 | 20667 | 1 |
| 484 | chi[3] | -0.10 | 0.03 | 5.59 | -6.91 | -2.06 | -0.73 | 1.15 | 10.28 | 38546 | 1 |
| 485 | chi[4] | -1.95 | 0.05 | 6.48 | -10.28 | -3.46 | -1.83 | -0.51 | 6.17 | 20327 | 1 |
| 486 | chi[5] | 1.02 | 0.04 | 5.80 | -3.89 | -1.16 | 0.04 | 1.91 | 11.55 | 25381 | 1 |
| 487 | chi[6] | -3.84 | 0.04 | 6.19 | -13.51 | -5.45 | -3.11 | -1.52 | 2.30 | 19263 | 1 |
| 488 | chi[7] | -1.98 | 0.04 | 5.88 | -10.62 | -3.23 | -1.86 | -0.72 | 6.93 | 27718 | 1 |
| 489 | chi[8] | -2.59 | 0.03 | 4.95 | -9.96 | -3.45 | -2.20 | -1.26 | 3.52 | 30293 | 1 |
| 490 | chi[9] | 1.44 | 0.03 | 5.26 | -3.52 | -0.73 | 0.49 | 2.37 | 12.07 | 29082 | 1 |
| 491 | chi[10] | -2.48 | 0.03 | 4.30 | -9.01 | -3.58 | -2.30 | -1.39 | 4.14 | 28444 | 1 |
| 492 | chi[11] | -2.12 | 0.03 | 6.17 | -11.88 | -3.62 | -1.78 | -0.33 | 6.41 | 32610 | 1 |
| 493 | chi[12] | 0.90 | 0.04 | 6.42 | -4.01 | -1.35 | -0.14 | 1.82 | 11.49 | 26833 | 1 |
| 494 | chi[13] | 0.01 | 0.03 | 5.85 | -6.33 | -1.94 | -0.63 | 1.35 | 10.10 | 34712 | 1 |
| 495 | chi[14] | -0.76 | 0.04 | 6.04 | -8.17 | -2.35 | -1.11 | 0.38 | 8.88 | 22827 | 1 |
| 496 | chi[15] | -0.19 | 0.07 | 8.43 | -6.31 | -2.09 | -0.82 | 0.92 | 9.83 | 13526 | 1 |
| 497 | chi[16] | -1.40 | 0.05 | 7.00 | -8.57 | -2.93 | -1.68 | -0.43 | 7.97 | 17139 | 1 |
| 498 | chi[17] | -1.97 | 0.03 | 5.61 | -11.42 | -3.37 | -1.75 | -0.41 | 6.49 | 37852 | 1 |
| 499 | chi[18] | -0.13 | 0.03 | 6.11 | -5.70 | -2.09 | -0.88 | 0.85 | 10.34 | 33396 | 1 |
| 500 | chi[19] | -0.86 | 0.08 | 8.93 | -9.55 | -2.51 | -1.08 | 0.58 | 9.14 | 14181 | 1 |
| 501 | chi[20] | -0.49 | 0.04 | 6.21 | -8.70 | -2.30 | -0.79 | 1.04 | 9.80 | 27651 | 1 |
| 502 | chi_0[1] | 2.44 | 0.05 | 6.92 | -1.26 | 1.47 | 2.33 | 3.36 | 7.07 | 18596 | 1 |
| 503 | chi_0[2] | 2.66 | 0.02 | 3.17 | -0.42 | 1.57 | 2.39 | 3.41 | 7.05 | 38581 | 1 |
| 504 | chi_0[3] | 2.71 | 0.02 | 3.16 | -0.67 | 1.63 | 2.48 | 3.54 | 7.30 | 40346 | 1 |
| 505 | chi_0[4] | 2.11 | 0.01 | 3.17 | -1.94 | 1.26 | 2.12 | 3.06 | 5.96 | 46298 | 1 |
| 506 | chi_0[5] | 2.81 | 0.03 | 4.71 | 0.00 | 1.53 | 2.36 | 3.45 | 7.63 | 19683 | 1 |
| 507 | chi_0[6] | 1.99 | 0.03 | 3.88 | -2.01 | 1.25 | 2.08 | 2.99 | 5.56 | 20733 | 1 |
| 508 | chi_0[7] | 3.06 | 0.03 | 4.53 | 0.74 | 1.84 | 2.61 | 3.62 | 7.46 | 18147 | 1 |
| 509 | chi_0[8] | 2.38 | 0.02 | 3.83 | -1.27 | 1.52 | 2.35 | 3.34 | 6.28 | 27529 | 1 |
| 510 | chi_0[9] | 2.96 | 0.02 | 3.74 | -0.45 | 1.71 | 2.59 | 3.74 | 8.17 | 32897 | 1 |
| 511 | chi_0[10] | 2.84 | 0.02 | 2.84 | 0.33 | 1.78 | 2.55 | 3.53 | 6.77 | 33529 | 1 |
| 512 | chi_0[11] | 2.61 | 0.02 | 3.17 | -0.31 | 1.53 | 2.34 | 3.34 | 6.81 | 30020 | 1 |
| 513 | chi_0[12] | 3.02 | 0.03 | 4.15 | -0.02 | 1.77 | 2.61 | 3.72 | 7.83 | 23182 | 1 |
| 514 | chi_0[13] | 2.44 | 0.02 | 3.87 | -2.03 | 1.42 | 2.33 | 3.41 | 6.88 | 35963 | 1 |
| 515 | chi_0[14] | 2.59 | 0.02 | 3.76 | -0.78 | 1.59 | 2.43 | 3.45 | 6.90 | 43349 | 1 |
| 516 | chi_0[15] | 2.52 | 0.05 | 6.45 | -1.55 | 1.48 | 2.36 | 3.40 | 6.93 | 15002 | 1 |
| 517 | chi_0[16] | 2.66 | 0.01 | 3.24 | -0.76 | 1.64 | 2.48 | 3.53 | 6.99 | 53450 | 1 |
| 518 | chi_0[17] | 2.16 | 0.02 | 3.26 | -1.95 | 1.31 | 2.19 | 3.18 | 6.16 | 40786 | 1 |
| 519 | chi_0[18] | 3.03 | 0.03 | 4.20 | 0.30 | 1.79 | 2.61 | 3.68 | 7.70 | 23585 | 1 |
| 520 | chi_0[19] | 2.43 | 0.02 | 3.83 | -1.58 | 1.47 | 2.34 | 3.37 | 6.81 | 36946 | 1 |
| 521 | chi_0[20] | 2.55 | 0.02 | 3.94 | -1.46 | 1.52 | 2.38 | 3.43 | 7.05 | 30406 | 1 |
| 522 | delta_0[1] | 1.11 | 0.00 | 0.14 | 0.85 | 1.02 | 1.11 | 1.20 | 1.39 | 38926 | 1 |
| 523 | delta_0[2] | 1.05 | 0.00 | 0.11 | 0.83 | 0.98 | 1.05 | 1.12 | 1.26 | 66376 | 1 |
| 524 | delta_0[3] | 0.84 | 0.00 | 0.11 | 0.61 | 0.76 | 0.84 | 0.91 | 1.06 | 53594 | 1 |
| 525 | delta_0[4] | 0.92 | 0.00 | 0.10 | 0.72 | 0.85 | 0.92 | 0.99 | 1.12 | 58054 | 1 |
| 526 | delta_0[5] | 0.97 | 0.00 | 0.12 | 0.72 | 0.90 | 0.97 | 1.04 | 1.17 | 46625 | 1 |
| 527 | delta_0[6] | 0.98 | 0.00 | 0.10 | 0.79 | 0.92 | 0.98 | 1.05 | 1.18 | 65330 | 1 |
| 528 | delta_0[7] | 1.07 | 0.00 | 0.10 | 0.86 | 0.99 | 1.06 | 1.13 | 1.27 | 67533 | 1 |
| 529 | delta_0[8] | 0.92 | 0.00 | 0.13 | 0.66 | 0.83 | 0.92 | 1.01 | 1.18 | 49771 | 1 |
| 530 | delta_0[9] | 0.80 | 0.00 | 0.15 | 0.52 | 0.70 | 0.79 | 0.89 | 1.10 | 35432 | 1 |
| 531 | delta_0[10] | 1.25 | 0.00 | 0.11 | 1.03 | 1.18 | 1.25 | 1.33 | 1.47 | 55695 | 1 |

| | | | | | | | | | | |
|---|---|---|---|---|---|---|---|---|---|---|
| delta_0[11] | 1.10 | 0.00 | 0.10 | 0.90 | 1.04 | 1.10 | 1.16 | 1.28 | 70168 | 1 |
| delta_0[12] | 0.92 | 0.00 | 0.10 | 0.73 | 0.85 | 0.92 | 0.98 | 1.11 | 71403 | 1 |
| delta_0[13] | 0.89 | 0.00 | 0.10 | 0.70 | 0.83 | 0.89 | 0.96 | 1.09 | 56016 | 1 |
| delta_0[14] | 1.21 | 0.00 | 0.10 | 1.02 | 1.14 | 1.21 | 1.27 | 1.40 | 74389 | 1 |
| delta_0[15] | 1.14 | 0.00 | 0.10 | 0.93 | 1.07 | 1.14 | 1.21 | 1.33 | 69998 | 1 |
| delta_0[16] | 1.08 | 0.00 | 0.11 | 0.87 | 1.01 | 1.08 | 1.16 | 1.30 | 63736 | 1 |
| delta_0[17] | 0.97 | 0.00 | 0.10 | 0.78 | 0.91 | 0.97 | 1.04 | 1.17 | 80936 | 1 |
| delta_0[18] | 1.12 | 0.00 | 0.10 | 0.94 | 1.06 | 1.12 | 1.19 | 1.32 | 67174 | 1 |
| delta_0[19] | 0.81 | 0.00 | 0.12 | 0.59 | 0.74 | 0.81 | 0.89 | 1.04 | 52180 | 1 |
| delta_0[20] | 1.19 | 0.00 | 0.14 | 0.91 | 1.10 | 1.19 | 1.28 | 1.47 | 54676 | 1 |
| eta[1] | 0.05 | 0.00 | 0.05 | -0.04 | 0.02 | 0.05 | 0.08 | 0.17 | 41428 | 1 |
| eta[2] | 0.02 | 0.00 | 0.04 | -0.06 | -0.01 | 0.02 | 0.05 | 0.11 | 71467 | 1 |
| eta[3] | 0.00 | 0.00 | 0.04 | -0.07 | -0.02 | 0.00 | 0.03 | 0.08 | 56256 | 1 |
| eta[4] | 0.01 | 0.00 | 0.06 | -0.13 | -0.03 | 0.01 | 0.05 | 0.14 | 58305 | 1 |
| eta[5] | -0.01 | 0.00 | 0.06 | -0.15 | -0.04 | 0.00 | 0.03 | 0.09 | 46133 | 1 |
| eta[6] | 0.04 | 0.00 | 0.07 | -0.09 | -0.01 | 0.03 | 0.07 | 0.22 | 49301 | 1 |
| eta[7] | 0.00 | 0.00 | 0.06 | -0.13 | -0.03 | 0.00 | 0.04 | 0.12 | 49441 | 1 |
| eta[8] | 0.02 | 0.00 | 0.07 | -0.11 | -0.02 | 0.02 | 0.06 | 0.17 | 52793 | 1 |
| eta[9] | -0.02 | 0.00 | 0.06 | -0.15 | -0.05 | -0.01 | 0.02 | 0.08 | 30824 | 1 |
| eta[10] | 0.01 | 0.00 | 0.07 | -0.15 | -0.03 | 0.01 | 0.05 | 0.13 | 40411 | 1 |
| eta[11] | 0.03 | 0.00 | 0.07 | -0.10 | -0.01 | 0.03 | 0.07 | 0.20 | 61125 | 1 |
| eta[12] | 0.02 | 0.00 | 0.06 | -0.09 | -0.01 | 0.02 | 0.06 | 0.16 | 67106 | 1 |
| eta[13] | 0.03 | 0.00 | 0.07 | -0.11 | -0.01 | 0.02 | 0.06 | 0.19 | 65854 | 1 |
| eta[14] | 0.03 | 0.00 | 0.06 | -0.10 | -0.01 | 0.02 | 0.06 | 0.16 | 54079 | 1 |
| eta[15] | 0.02 | 0.00 | 0.07 | -0.12 | -0.02 | 0.02 | 0.05 | 0.16 | 73922 | 1 |
| eta[16] | -0.01 | 0.00 | 0.06 | -0.14 | -0.05 | -0.01 | 0.03 | 0.09 | 43406 | 1 |
| eta[17] | 0.03 | 0.00 | 0.06 | -0.08 | -0.01 | 0.03 | 0.06 | 0.16 | 60087 | 1 |
| eta[18] | 0.01 | 0.00 | 0.06 | -0.12 | -0.02 | 0.01 | 0.05 | 0.15 | 73311 | 1 |
| eta[19] | 0.00 | 0.00 | 0.06 | -0.15 | -0.04 | 0.00 | 0.04 | 0.11 | 42933 | 1 |
| eta[20] | 0.03 | 0.00 | 0.04 | -0.06 | 0.00 | 0.03 | 0.05 | 0.11 | 55805 | 1 |
| delta_0_av | 1.00 | 0.00 | 0.19 | 0.61 | 0.88 | 1.00 | 1.12 | 1.38 | 80947 | 1 |
| eta_av | 0.01 | 0.00 | 0.07 | -0.13 | -0.02 | 0.01 | 0.05 | 0.16 | 70943 | 1 |
| chi_av | -0.90 | 0.02 | 6.14 | -9.46 | -2.62 | -1.14 | 0.59 | 9.09 | 100000 | 1 |
| chi_0_av | 2.48 | 0.02 | 4.49 | -1.54 | 1.48 | 2.36 | 3.39 | 6.87 | 60077 | 1 |
| lp__ | 485.14 | 0.06 | 10.18 | 464.37 | 478.46 | 485.47 | 492.15 | 504.15 | 25442 | 1 |

Samples were drawn using NUTS(diag_e) at Thu Oct 26 16:10:03 2017. For each parameter, n_eff is a crude measure of effective sample size, and Rhat is the potential scale reduction factor on split chains (at convergence, Rhat=1).
